# TS-Zarr: a cloud-optimized Zarr layout for viewing and analyzing biomedical time series

**DOI:** 10.64898/2026.11.16.767094

**Authors:** Rohan Shah, Sumita Garai, D.D. De Freitas, Van QT Truong, Wolfgang Ganglberger, Tucker Paron, M. Brandon Westover, Nishant Sinha, Joost Wagenaar

## Abstract

**Background:** Biomedical time series are recorded at high sampling rates for hours to days across tens to hundreds of channels. Each recording occupies gigabytes to terabytes of storage. FAIR data sharing requirements often place recordings in cloud storage. Visual analysis requires downsampling windows wider than a few seconds to the pixel width of the screen. Visualization tools therefore either read a local copy of the recording or query a dedicated resampling server in front of the data.

**Findings:** We present TS-Zarr, a Zarr v3 layout for time series data. It stores the samples of each channel alongside a pyramid of summary statistics at progressively coarser levels. Zarr on cloud storage offers low-latency, high-throughput parallel reads. The layout lets a reader fetch only the channels and windows it needs. These targeted reads scale better than the same reads from HDF5 and EDF. Precomputed levels trade additional storage and error-bound signal transformation for faster and more scalable visual analysis that moves from the whole recording to narrow windows. As users grow and windows widen, precomputed reads from cloud storage scale beyond the point at which a resampling server fails.

**Conclusions:** Precomputed levels let researchers visualize a recording without downloading a local copy or querying a dedicated resampling server. The time to read a view remains bounded as the window under view widens. Zarr on cloud storage also provides efficient, highly parallelizable, targeted reads of raw sample data for analysis.

## Background

### Visualizing time series at scale

Visual analysis is widely used to explore time series data, and line charts have been shown to be effective for most analysis tasks [1, 2]. Visualizing a time series lets researchers form questions before they analyze specific segments of the data [3, 4]. This pattern is summarized as overview first, zoom and filter, then details on demand [5]. As a series grows, a chart of the whole series draws many samples in each column of pixels on the screen. Rendering every sample is slow and hides events shorter than a pixel column [6]. Downsampling addresses this by drawing a reduced set of values for each column. Since the downsampled series serves human observation, only the values that the eye can resolve need to be kept [6]. For a line chart these are the minimum and maximum of each column [7].

Biomedical recordings run for hours to days across tens to hundreds of channels. Scalp elec-troencephalography (EEG) records tens of electrodes at a few hundred samples per second [8, 9], intracranial EEG records hundreds of contacts at kilohertz rates, and silicon probes (Neuropixels) record hundreds of sites at 30 kHz [10]. Polysomnography adds respiratory, muscle, eye, and oximetry channels at mixed rates over a night, and epilepsy monitoring continues for days. Clinical review of EEG is paged, at 10 s per screen [9]. A day-long recording can span thousands of pages. Figure 1 shows one channel of a synthetic 2 kHz recording at three window widths, from a 10 s page to 0.1 s.

**Figure 1:**
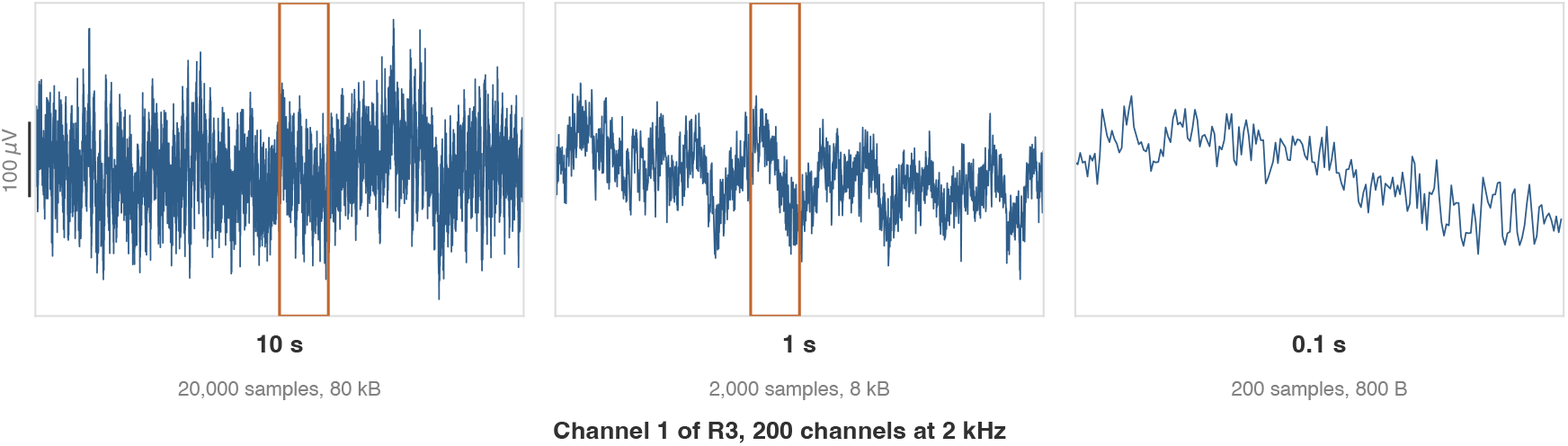
Channel 1 of the synthetic recording R3 (Appendix C) at windows of 10 s, 1 s, and 0.1 s. The orange box marks the window of the next panel.

In current systems the downsampling is computed when a view is requested, on a resampling server or on the user’s machine. Relational databases run the downsampling as a query [7], and time-series databases offer it as a built-in function [11]. Plotting libraries run it in a callback behind the figure [12], and web services cache the downsampled result (as values or as rendered images) and send it to the client [2, 7]. In each design the resampling server reads the samples under the requested window, downsamples them to the width of the screen, and returns the result (Figure 2). Because the work is repeated for every window and every user, the load grows with the number of users. In the example of Jugel et al., 100 engineers who each view 12 h of 100 Hz data request 432 million rows in total [7]. Caching the downsampled results [13] and keeping extrema at several levels inside the database [14] reduce the repeated work, but the downsampling remains inside the service.

**Figure 2:**
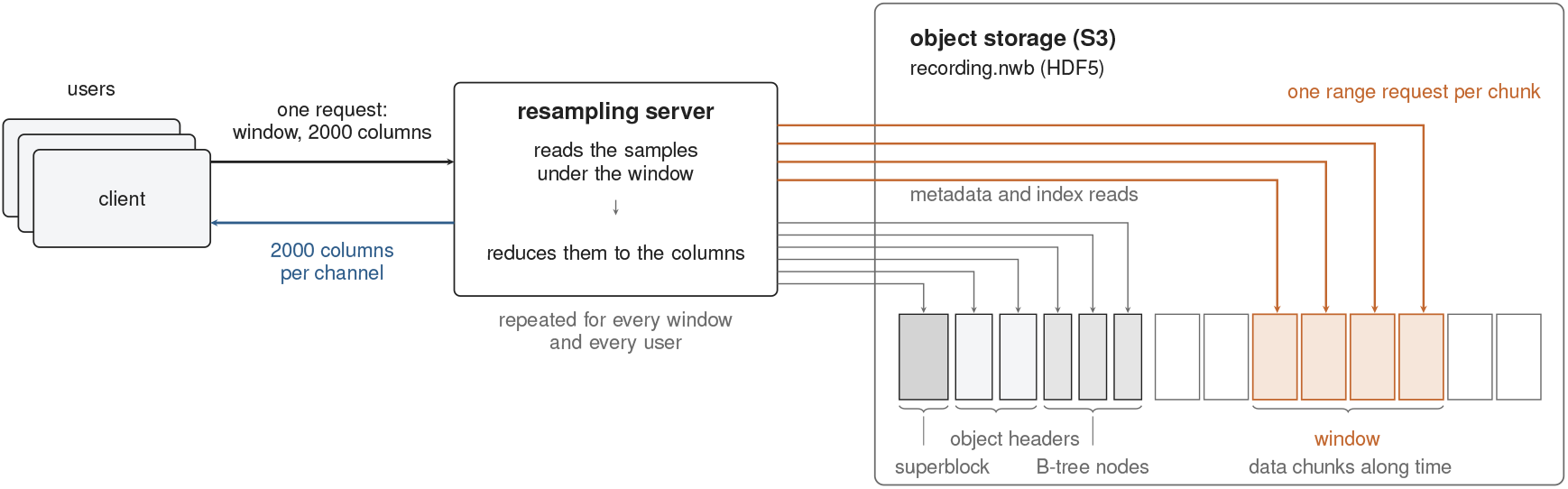
Downsampling at query time. Each client sends one request for a window to a resampling server beside the data. The server reads the file index and the chunks under the window by range request, downsamples the window to the width of the screen, and returns the downsampled series to the client.

Locally installed tools such as EDFbrowser, EEGLAB, and MNE-Python draw a recording from a copy on the user’s machine [15–17]. Each user therefore needs to store a copy of the recording and update it if the source changes.

Downsampling at query time is practical for a time series that is still growing. Systems such as M4 and time-series databases were built for this use case [7]. Real-time acquisition in biomedicine has this requirement when signals are viewed as they are recorded, for example at the bedside. Such signals are often stored in databases or other appendable formats that support query-time downsampling.

This paper is concerned with recordings whose acquisition is complete, for instance a polysomnogram after the night or an EEG after an admission. Such recordings are typically distributed as static files, in EDF+, NWB, or proprietary formats [18, 19], and published to archives for reuse. They are rarely modified, but they are read for years in clinical review, in teaching, and in retrospective analysis. The downsampling can therefore be computed once, when the recording is deposited, and stored where any reader can fetch only the channels and windows it needs, without a local copy or a server.

Stored summaries at several levels are established in other fields. Genome browsers keep the count, minimum, maximum, sum, and sum of squares of each bin at several zoom levels in BigWig files and read them over HTTP by range request [20]. Monitoring databases have kept rollups of this kind for decades [21, 22], and binned summaries computed in advance are the standard approach to interactive views of large point data [23, 24]. Existing file formats for biomedical time series have not adopted these techniques, an absence that may reflect the focus on real-time acquisition and the dependence of clinical review on filtered and montaged views.

### Zarr for biomedical time series

FAIR data sharing has led researchers to deposit biomedical data in cloud object storage [25]. Scientific formats designed before object storage was common, such as HDF5, “perform numerous small seek / read operations when opening files and reading data”, which makes them slow to read from an object store [26]. Zarr was designed for parallel and distributed computation on large arrays [27]. It stores each chunk of a compressed N-dimensional array under its own key, with the array metadata in JSON beside the chunks [27, 28]. Readers can fetch only the chunks they need from object storage, independently and in parallel, at scale [29]. In the Pangeo benchmark, NetCDF4/HDF5 read speeds on object storage were “an order of magnitude slower than Zarr” [26].

Other biomedical data formats have adopted Zarr for cloud-optimized storage. OME-Zarr stores microscopy images in Zarr as chunked multiscale pyramids [30], and VCF Zarr stores genomic variants in Zarr and reports large gains in storage and analysis cost over VCF at biobank scale [31]. Zarr has readers in Python, Java, C++, JavaScript, and Rust and holds datasets in geoscience, genomics, and bioimaging [26, 30, 31].

OME-Zarr was designed to complement existing bioimaging formats such as OME-TIFF and HDF5 [32]. It precomputes a multiscale pyramid [30], which lets a reader fetch a zoom level more efficiently than on-the-fly translation, which “must be repeated on each use” [32], similar to the query-time downsampling of time series described above.

Existing time series formats already allow random reads from object storage. EDF+ and MEF keep a whole biomedical time series recording in one file, with a header and a fixed data layout that locate any channel and time span [18, 33]. NWB stores it in HDF5, whose chunk index serves the same purpose [34, 35]. A reader can fetch the byte ranges it needs from such a file in object storage by range request [36–38]. However, Zarr serves the same reads better, since each chunk is a separate object that many readers can fetch in parallel. For example, NWB on Zarr, the hdmf-zarr backend, writes the NWB data model to Zarr and reads faster from cloud storage than either alternative (Analysis) [39, 40].

These formats typically store samples along a time axis, placing every channel’s samples for a span of time contiguously. This layout is efficient for reads of every channel in a fixed window and less efficient for reads of a single channel or a subset of channels. Because they store only the recorded samples, with no precomputed downsampling or summary statistics, a wide view of a long recording reads a large volume of sample data over the network.

We present TS-Zarr, a Zarr v3 layout for biomedical time series recordings. It stores the recorded samples of every channel alongside summary statistics at progressively coarser levels. Each level summarizes the samples in consecutive bins of time. For each view, a visualization tool opens a recording over HTTP by range request and reads the level whose bins are no wider than a pixel column. For 200 intracranial channels at 2 kHz, a view of a whole 24 h recording reads 4.2 MB in place of 138 GB (Table 1).

**Table 1:** A viewer 2000 pixels wide reads the sizes below at seven window widths of a 24 h recording of 200 channels at 2 kHz. Sizes are before compression. A bin is 8 bytes, 4 for the minimum and 4 for the maximum.

| window | one pixel column | level read | bins or samples per channel | size read, 200 channels | size of the samples under the window |
|---|---|---|---|---|---|
| 0.1 s | 50 *μ*s | raw | 200 samples | 0.16 MB | 0.16 MB |
| 1 s | 0.5 ms | raw | 2000 samples | 1.6 MB | 1.6 MB |
| 10 s page | 5 ms | 1 (2 ms bins) | 5000 | 8 MB | 16 MB |
| 1 min | 30 ms | 2 (8 ms bins) | 7500 | 12 MB | 96 MB |
| 10 min | 300 ms | 4 (128 ms bins) | 4688 | 7.5 MB | 960 MB |
| 1 h | 1.8 s | 5 (512 ms bins) | 7032 | 11 MB | 5.76 GB |
| 24 h | 43.2 s | 8 (32.8 s bins) | 2637 | 4.2 MB | 138 GB |

Like OME-Zarr, this format is designed to complement existing formats such as EDF+ and NWB. We (i) describe the layout and how it supports efficient and accurate visualization of time series, (ii) measure its read time against EDF and NWB, (iii) compare how it scales with a resampling server, and (iv) discuss its tradeoffs.

## Format Design

### Layout

A TS-Zarr bundle represents a single recording of biomedical time series data. Time-aligned formats such as EDF and NWB typically store the samples of every channel together along a time axis. This type of layout requires special handling for channels of different sample rates, and a read fetches a window of samples across every channel. We instead store each channel in its own Zarr group, a channel-aligned layout (Figure 3), which allows each channel to maintain its own sample rate and a reader to fetch only the channels it needs, in parallel.

**Figure 3:**
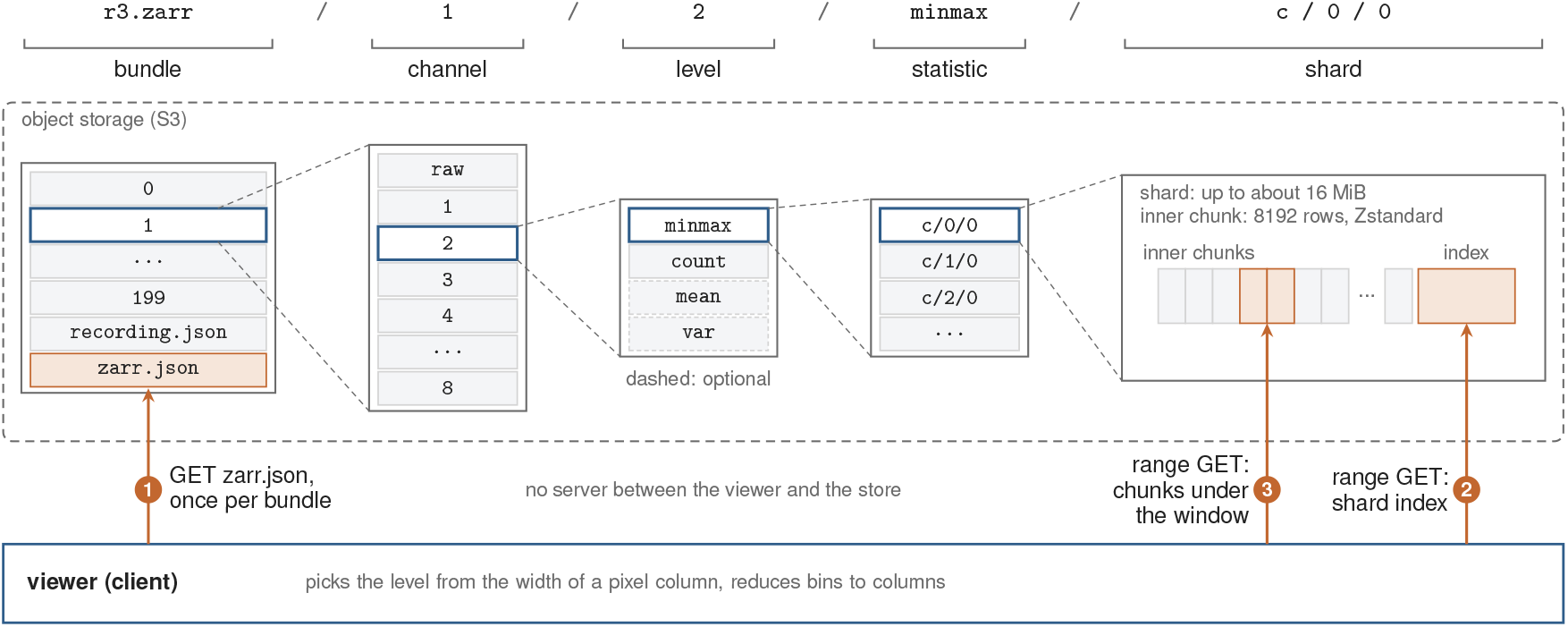
A viewer reads one view of a TS-Zarr bundle in object storage with two range requests per channel. Each object key is a path of channel, level, statistic, and shard. The viewer reads the root metadata once and chooses the level from the width of a pixel column. It then fetches the shard index and the inner chunks under its window. It reads from the store directly, with no server in between.

A continuous channel stores its samples in an array named `raw`. Like every array in the bundle, it is chunked along time, sharded, and compressed (Appendix A), and a viewer reads the chunks under its window by range request.

Alongside `raw`, a continuous channel stores a pyramid of downsampled levels numbered 1 to *k* (Figure 4). Each level is coarser than the one below it by a configurable step. At a step of four, level *k* holds one bin for every 4^*k*^ samples. A level stores two statistics of the finite samples in each bin, `minmax` and count, and may add `mean` and `var` for the views that need them.

**Figure 4:**
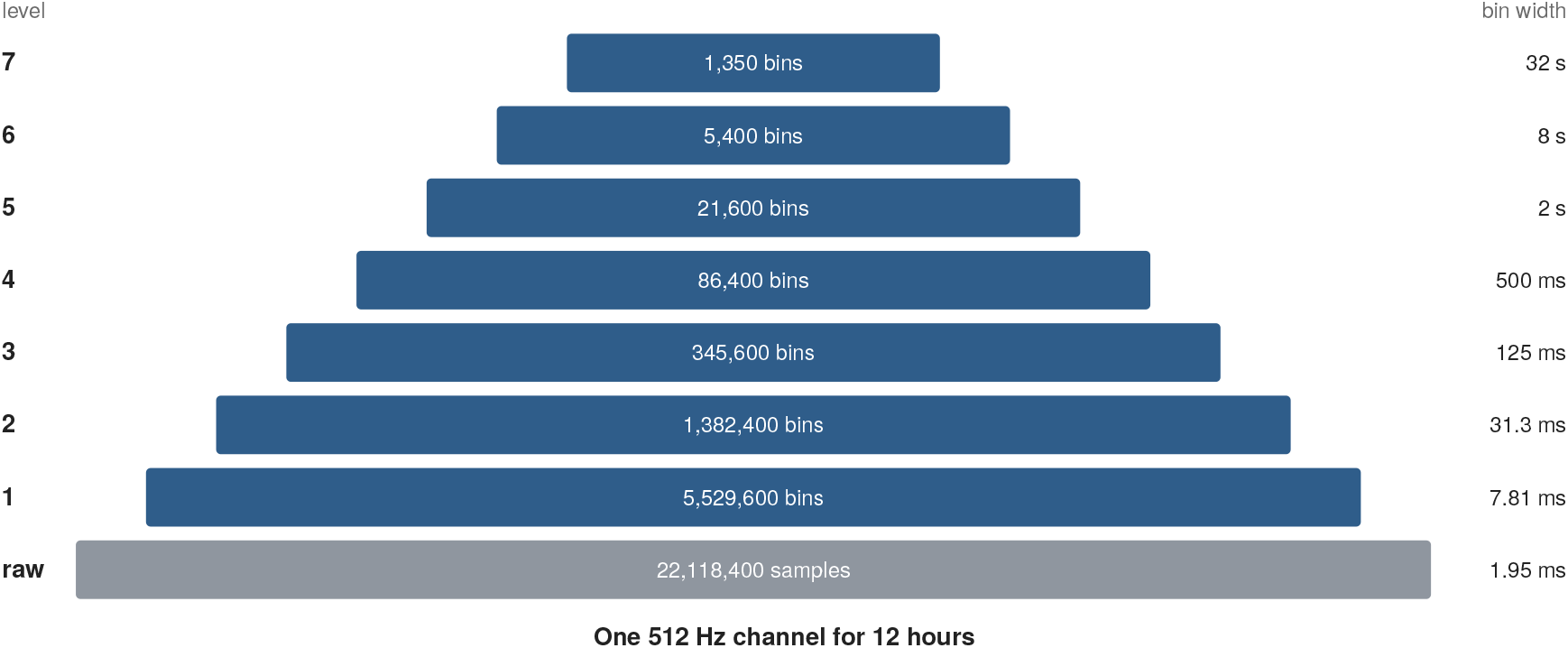
At a step of four, level *k* of a channel of 12 h at 512 Hz holds one bin for every 4^*k*^ samples. The length of each bar is the logarithm of the number of bins in the level. The pyramid ends before a level would hold fewer than 1024 bins.

An event channel stores the spike times of sorted units or the annotations of a recording, such as seizures and sleep stages, as parallel arrays indexed by event. Because the event times are kept in order, a binary search can find the events in a given window. An upper bound on event duration lets the search also return intervals that began before the window. Optional arrays hold a label, a value, text, the channels an event applies to, and a waveform. A dense event channel carries levels of event counts for each label. A firing-rate raster at overview width reads bin counts instead of every spike (Figure S1, Appendix A).

### Levels for visualization

OME-Zarr stores an image as a pyramid of lower resolutions. A viewer can open a 1 TB plate as a 2560 by 1822 pixel image [30]. Similarly, the amount of data a viewer needs to display a window of a time series depends on the width of the screen and remains bounded as the length of the window increases. A viewer reads the coarsest level whose bins span no more time than one pixel column. If even the finest level is too coarse, the viewer reads the original samples (`raw`). At a step of four, the chosen level has between one and four bins per pixel column. A chart 2000 pixels wide reads between 2000 and 8000 bins per channel. Table 1 shows what a viewer reads from a 24 h recording of 200 intracranial channels at 2 kHz, which is 138 GB of samples in float32. A window of 1 s or less spans at most 2000 samples per channel and is read from `raw`. A 10 s window, the width of a review page, reads 8 MB from level 1 in place of 16 MB of raw samples. Across windows from 10 s to 24 h, the read stays between 4.2 and 12 MB while the raw samples under the window grow from 16 MB to 138 GB. A window of one week would read 7.4 MB.

### Extrema for line charts

A line chart of a time series draws a straight line from each sample to the next. Downsampling a window to fit its pixel columns distorts how that line moves. When a pixel column spans several samples, the lines drawn in that column are combined together into a vertical bar from the lowest sample to the highest. The line enters the column at the height of its earliest sample and leaves at the height of its latest. Jugel et al. show that the minimum, maximum, earliest, and latest sample of each column are required to reproduce a two-color line chart (the line in one color on a background of another) without error [7]. Averaging within a column (piecewise aggregate approximation) acts as a low-pass filter and attenuates the extrema, and the median, medoid, and mode distort the rendered line in the same way [7]. The methods that preserve the shape select samples instead. The MinMax method uses the minimum and maximum of each column. Its vertical bars show how far the signal moves in each column. They do not trace a single path through the signal as a single point per column does, but they keep the extrema that averaging attenuates.

Each level of a continuous channel stores minmax, the minimum and maximum of the finite samples in each bin. The minimum of the minima of several bins is the minimum of their samples, and the same holds for the maxima. A level can therefore be derived from the level directly below it and have the same minima and maxima as a level computed from the raw samples. A bin stores NaN only when it contains no finite sample, such as when a gap, a stretch of non-finite samples, covers a whole bin. A shorter gap is recorded as a lower count for the bin.

### Statistics for derived views

Clinical review often uses filters to focus on particular frequencies and montages to display signals relative to a chosen reference. Common examples are bipolar or average-reference montages with a 0.5 to 70 Hz bandpass, a filter suited to each channel class in polysomnography, and trends of power over hours. The format lets a producer store the `mean` and `var` of each bin beside the required `minmax` and `count`. A viewer can compute these views from a level that stores them. Because montages are linear functions of the samples, they can be computed from the `mean` of each bin. The mean of a montaged signal over a bin equals the montage of the channel means over that bin (Appendix B). A viewer can therefore apply a montage to the channel means of any level as it would to the samples.

Applying a linear filter to the `mean` of a level and not to the samples comes with some error. A level can represent frequencies only up to its Nyquist frequency, or half its sampling rate, since it has one value per bin. The mean of a bin attenuates content above that frequency without removing it, and the remainder appears at a lower frequency (aliasing). The filter must be redesigned for the rate of the level, with a passband below the Nyquist frequency of the level. For the clinical bandpass of 0.5 to 70 Hz on a 2 kHz channel, level 1 has a rate of 500 Hz and a Nyquist frequency of 250 Hz, above the 70 Hz edge. The bandpass cannot be applied to level 2, whose Nyquist frequency of 62.5 Hz is below the edge. In Figure 5, the bandpass on level 1 differs from the bandpass on the samples by a root mean square of 0.98 *µ*V, against 52 *µ*V for the montage. Most of the difference comes from content within 70 Hz of 500 Hz. The mean of four samples attenuates that content without removing it, and it appears in the passband. The mean also scales down anything shorter than a bin by the fraction of the bin that it fills. At overview widths a viewer can draw it as the trend inside the `minmax` bars.

**Figure 5:**
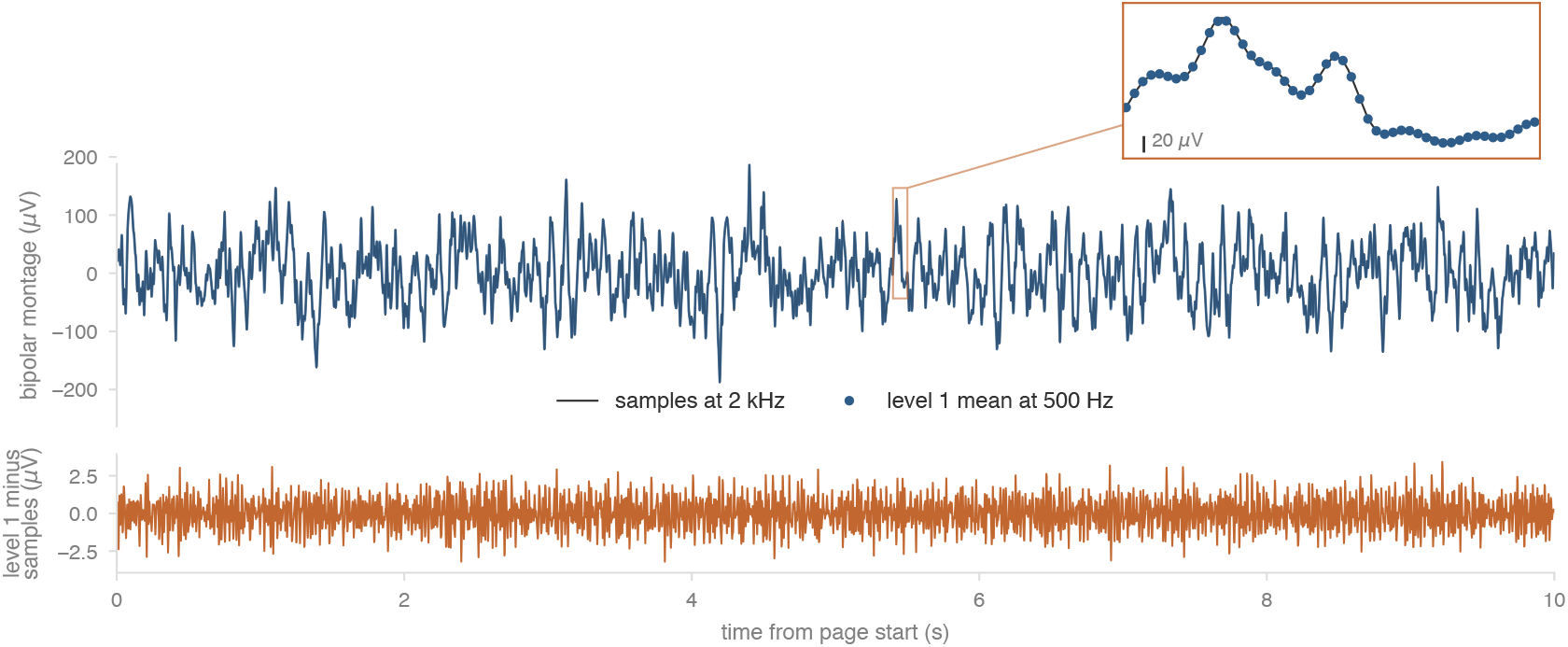
A bipolar montage of channels 1 and 2 of the synthetic recording R3 over the 10 s page of Figure 1 is bandpass filtered from 0.5 to 70 Hz with a fourth-order Butterworth filter, applied forward and backward. The filter is applied to the samples at 2 kHz and to the `mean` of level 1 at 500 Hz. The upper panel overlays the two filtered signals. Its inset enlarges the 0.1 s window of Figure 1, with the samples as a line and the bin centers of level 1 as dots. The lower panel shows the filtered signal from level 1 minus the filtered signal from the samples at the bin centers of level 1. The difference has a root mean square of 0.98 *µ*V and a largest magnitude of 3.4 *µ*V, against a root mean square of 52 *µ*V for the montage from the samples.

A trend panel plots power over hours, which the mean of a bin does not carry. The amplitude of an EEG across a night, the chin EMG tone that falls in REM sleep, and the root mean square (RMS) of any channel can be computed from `var`, the variance of each bin, together with mean. A viewer can read var at the coarsest level for every channel in one small request to see which channels were flat, saturated, or noisy.

Coverage and gaps can be read from `count`, the number of finite samples in each bin, which the format requires alongside `minmax`. A bin with part of a gap still has a minimum, a maximum, and a mean from its finite samples, and only its `count` falls below the bin width. A count of zero means a gap spans the entire bin. A count one less than the bin width means one sample is non-finite, which a viewer can find in a finer level or in the samples. A viewer can divide count at the coarsest level by the bin width to get the recorded fraction of every bin. A coverage strip for a week requires only one request.

Views that cannot be computed from the bin statistics require reading `raw`. These include nonlinear operations such as a median filter or artifact rejection, montages across channels of different rates, and any view that needs content above the Nyquist frequency of a level.

## Analysis

We show where Zarr outperforms HDF5 and EDF and how precomputed levels scale read latency and throughput for wide windows and for many users, where a resampling server slows until failure.

### Zarr against HDF5 and EDF

We measured how the time to read the raw samples of a window from Zarr compares with HDF5 and EDF. Two of the four formats are Zarr, NWB on Zarr and TS-Zarr, and the others are NWB on HDF5 and EDF. The benchmark read only the raw arrays of TS-Zarr. Its precomputed levels did not influence this comparison. The design follows the latency benchmark of OME-NGFF [32], with time windows in place of chunks. We wrote four synthetic recordings (Table S1) in these formats, NWB twice, once chunked along time and once chunked by channel. We read windows of 1 s to 10 min from one channel up to every channel, from local disk, behind an HTTP server, and from S3. Figure 6 shows the reads of R3 from S3. Appendix C describes the methods and reports the results for each recording and store.

**Figure 6:**
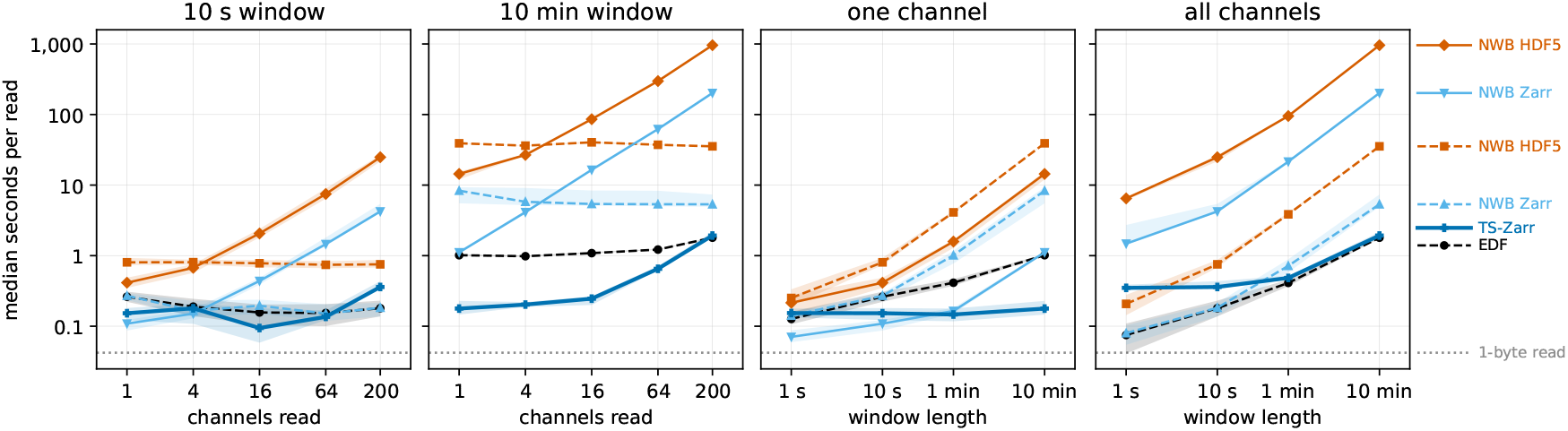
Median seconds per read of the raw samples of the first hour of R3 from S3, on log axes. The two left panels read windows of 10 s and 10 min from a growing number of channels. The two right panels read one channel and all 200 channels at growing window lengths. Solid lines are formats that store each channel separately (TS-Zarr and the NWB files chunked by channel). Dashed lines are formats that store a span of time of every channel together (EDF and the NWB files chunked along time). A TS-Zarr read fetches the raw arrays and no level. Bands span the 25th to 75th percentile of the reads, and the dotted line is the median of a 1-byte read from the same store. The medians cover 4,535 reads. Each combination of format, read, and window has between one and 100 reads, and it stops after the read that brings its total read time to at least 60 s.

The time to read a window of a time series from object storage depends on the number of requests and on how many of them can be made concurrently. An HDF5 file is stored as one object. The HDF5 reader used in our benchmark fetched the requested chunks sequentially. A 10 min read of one channel of NWB on HDF5 chunked along time, for example, made 935 requests and took 39 s. Zarr stores each chunk or shard as its own object, which a reader can request concurrently. On R3 and R4 from S3, NWB on Zarr read every window faster than NWB on HDF5 with the same chunk shape, by 1.8 to 23 times. Similarly, TS-Zarr was at least 1.4 times faster than NWB on HDF5 over every window, from reads of one channel to reads of many channels.

For short windows over a single channel, every format reads in a fraction of a second. As the window grows, however, the alignment of the data determines which reads stay fast. EDF and the NWB files chunked along time store every channel of a span together, and a read of one channel necessarily transfers all. TS-Zarr stores each channel apart, and a read of one channel touches only its own objects. It read one channel of R3 in two requests and under 0.2 s at every window from 1 s to 10 min, which at 10 min was 81 times faster than NWB on HDF5 chunked by channel. For blocks of 16 and 64 channels, a fraction of the channels of R3 and R4, TS-Zarr was the fastest format at every window tested from S3, by 1.1 to 15 times over the next fastest.

Stored as samples alone, every format occupied about half the size of the first hour of R3 as float32, with EDF at 16 bits and the others compressed (Table 2). Grouping Zarr chunks into shards reduces the number of objects. For example, NWB on Zarr chunked by channel wrote 175,891 objects without shards and 491 with them.

**Table 2:**
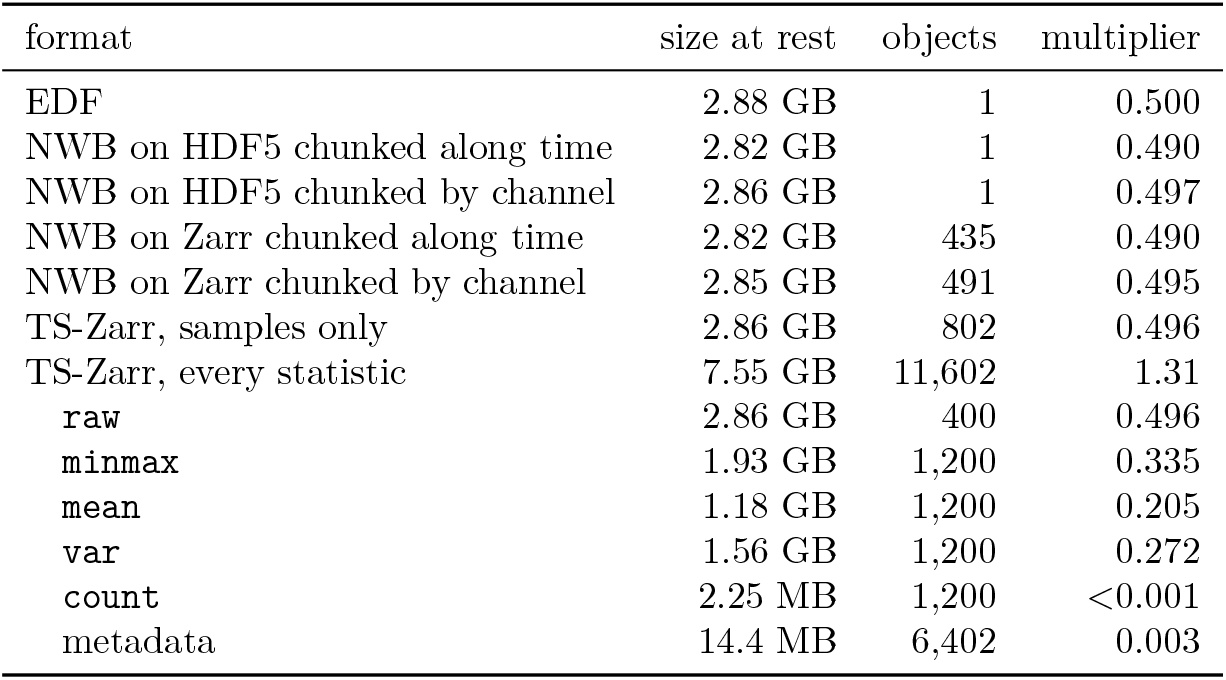
Size at rest and number of objects for the first hour of R3 in each format. The multiplier is the size divided by the 5.76 GB of the samples as float32. The indented rows divide the full TS-Zarr row by member. Appendix C (Table S2) reports every recording.

### Precomputed levels against on-demand resampling

We measured how reading the precomputed levels of TS-Zarr compares with resampling the samples on demand to visualize a time series. A resampling server read the samples under the window from the raw arrays in S3 and downsampled them to 2000 pixel columns on each request, without a cache. For the precomputed read, a user’s machine fetched the `minmax` of a level from S3 and downsampled its bins to fit the same 2000 pixel columns. Windows ran from 10 s to 24 h of recording R3, with 1, 10, 100, and 1000 simulated users running concurrently. Each user sent its next request when its previous request completed. Figure 7 shows the result.

**Figure 7:**
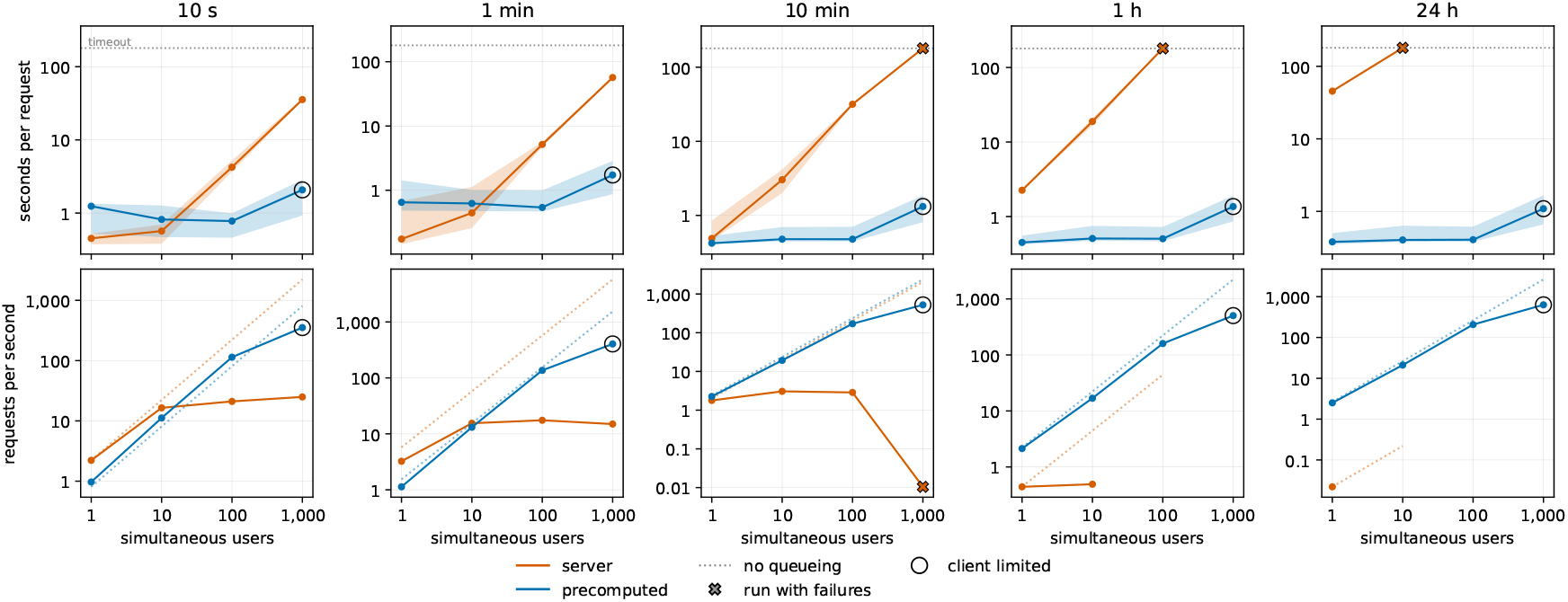
The resampling benchmark on R3 against the number of users. Median seconds per request with a band from the 5th to the 95th percentile (top) and requests completed per second (bottom). Dotted lines show the rate if every request took the 1-user median. Crosses mark runs with failures and rings mark client-limited runs (Appendix C).

The response time of a resampling server grows with the window, since it reads all samples within it, while a precomputed read depends only on a fixed number of pixel columns. With one user and small windows of 10 s and 1 min, the resampling server performed without issue, since it read little. However, it took 2.3 s per request at 1 h and 45 s at 24 h, where it transferred 69 GB of compressed samples from S3 at close to the 12.5 Gbps network of the benchmark instance. In contrast, the precomputed read took 0.38 to 1.24 s at every window from 10 s to 24 h, and at 24 h it read level 8 in 200 requests and 2.0 MB, which is 120 times faster than the server (Appendix C, Table S3).

The resampling server also shares a fixed set of processors between its users. As users were added, its CPU use rose to 68 to 93% of its 32 vCPU and its throughput leveled off. The median request time grew to 32 s at 10 min with 100 users. Requests failed at the 180 s timeout with 10 users at 24 h. The precomputed read moves the downsampling to the machine of each user and reads objects from S3. Its median request time stayed under 0.82 s up to 100 users across windows while its throughput grew with the users. At 1000 users the median rose to 1.1 to 2.1 s, but those runs put more users on each load generator than its calibrated limit, and S3 handled them without a failure. On R4 with 100 users, resampling took 19 s at 10 s and failed at 10 min, while the precomputed read took under 1 s.

## Discussion

Designing a data format such as TS-Zarr involves tradeoffs. Precomputed levels give a steady low latency and a high throughput from cloud storage, but they cost storage. Storing statistics in place of samples means some views cannot be computed from a level. Storing each channel apart favors reads of a few channels over reads of every channel. This section discusses each tradeoff and how a producer can configure the format to reduce some of the downsides. It then describes the downsampling methods that a level supports.

### Storage cost of levels

The largest cost of precomputing a pyramid of levels is the additional storage they need. A producer can tune that cost by choosing the step between levels, the number of levels, and the statistics to store.

At a step of four, level 1 has one bin for every four samples, and each later level has a quarter as many bins as the one before it. All levels together have one bin for every three samples 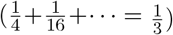. At a step of two they have one bin per sample. A bin has two values in `minmax` and one in each of `count`, `mean`, and `var`. The required minmax and count therefore add as many values as there are samples, and each of mean and var adds a further third.

Compression significantly helps reduce the amount of extra storage required to store the pyramid. In the first hour of R3, `minmax` was 0.68 of the size of the compressed samples, `mean 0.41`, var 0.55, and count less than 0.1% (Table 2). Counts of complete bins are constant and compress to almost nothing. The required statistics added about two thirds of the size of the samples, and all four added 1.6 times that size.

The step trades storage against read size. A smaller step stores more levels and more values, and a viewer reads fewer bins per pixel column. A larger step does the opposite. With a step of four and all four statistics, level 1 holds five values for every four samples. That is more than the samples themselves, and it describes the signal in less detail. Wherever level 1 could serve a request, reading raw costs less. With only the required statistics, level 1 holds three values for every four samples, a saving of one quarter. Level 1 is the largest level and the closest in size to the samples, so most of the storage that adds little is in level 1. The specification therefore lets the step vary by level. A producer can start the pyramid at a larger step, or leave `mean` and `var` out of the finest levels (both are optional members of each level).

The specification sets no maximum number of levels. The reference writer stops adding levels when the next would have fewer than 1024 bins.

### Limits of the stored statistics

Precomputed statistics reduce samples to a few numbers. A view computed from these statistics cannot always match the same view computed from the samples they cover. The differences take two forms. The statistics of a combination of channels, such as a montage, are not determined by the statistics of each channel. The mean of a bin also loses oscillations that are fast relative to the bin.

A montage drawn from `mean` is exact for the mean of each bin, but it attenuates the spikes that a montage of the samples carries. The minimum and maximum of a montage cannot be computed from the minimum and maximum of its channels (Appendix B). A viewer therefore cannot communicate the range that a montage of the samples would.

Mean statistics cannot represent oscillations faster than half the rate at which they are stored. Level 1 of a 2 kHz channel stores one mean for every four samples, or 500 values per second. It cannot represent oscillations faster than 250 Hz. Averaging four samples reduces a 320 Hz fast ripple to 0.47 of its amplitude and moves it to 180 Hz, inside the ripple band. However, a notch filter for the 50 or 60 Hz line frequency can be applied to level 1 as it would to the samples, since both frequencies are well below the limit.

### Sample alignment

A time-aligned layout stores the samples of every channel together for each span of time, and a channel-aligned layout stores each channel apart. Each layout comes with tradeoffs.

Time alignment pays off when a read spans every channel, since the samples can be read contiguously with little overhead. However, storing channels with mixed sampling rates requires special handling, often a separate grouping, which reduces the benefits of time alignment and makes the format more complex.

Channel alignment pays off when a read covers only some of the channels, or when channels are read independently, for example by different processes. It also allows each channel to carry its own sampling rate without special handling.

We chose channel alignment, which makes the format simpler, over optimizing reads that cover every channel. Offering time alignment is an option for a future version of TS-Zarr, to target recording-wide analysis.

### Downsampling methods

minmax supports drawing a vertical bar from the minimum to the maximum of each pixel column. The extrema that averaging attenuates remain visible. However, the bars form a dense view of the time series. They show how far the signal moves in each column but not the path it takes, and they make the shape of the time series harder to follow. Other methods for visualizing time series select samples that follow the line chart of the samples more closely.

M4 selects the minimum, maximum, earliest, and latest sample of each column in order to reproduce a two-color line chart [7]. Without the earliest or latest sample, M4 cannot be fully reproduced from a level. However, the error is confined to the pixels where the line joining two columns extends past the end of a bar.

The largest-triangle-three-buckets method (LTTB) selects the one sample per bucket that forms the largest triangle with its neighbors, which gives an effective visual representation of the original series [6]. It makes a sequential pass over all samples. MinMaxLTTB first preselects the minimum and maximum of each sub-bucket and then applies LTTB to the preselected points. The LTTB step scales with the output size, and the MinMax pass is linear in the number of samples [2].

Using `minmax`, a viewer can compute a variant of MinMaxLTTB by applying LTTB to the bins of the coarsest level that has at least two bins per pixel column. At a step of four, that gives between four and sixteen preselected points per column. Van Der Donckt et al. report that a preselection ratio of four or six “already results in high visual similarity to LTTB” [2]. A viewer that places each extremum at the center of its bin displaces it by at most half a bin, which is at most a quarter of a pixel column (Appendix B). We have not scored this variant against the exact algorithm on the templates of Van Der Donckt et al. [2].

## Conclusions

TS-Zarr complements the existing formats in which biomedical time series recordings are commonly archived and exchanged. EDF provides a simple, efficient model for storing time series that has served clinical exchange since 1992 [18, 41]. NWB improves on formats such as EDF for neuro-physiology. It has a rich data model, schema, and validator, and a large ecosystem of tools for reading, analyzing, and sharing data across many modalities [19, 35]. BIDS organizes recordings into datasets with standard names and metadata for sharing and reuse [42, 43]. TS-Zarr adds a layout optimized for object storage, which allows collaborative viewing and analysis of time series recordings at scale. It removes the need for users to download local copies of ever larger files and for data owners to run resampling servers that users need to query.

Our analysis shows that Zarr outperforms HDF5 and EDF in reading targeted windows and channels from cloud storage. Precomputation allows a viewer to read any window of a recording in a time set by the width of the screen rather than by the size of the window, from a 10 s page to a 24 h overview. Because reads scale with cloud storage, a recording can be served to millions of users, without the throughput limit of a resampling server. The statistics minmax, mean, and var allow a viewer to draw a time series at any width in a form that supports common visual analysis techniques, such as montages, filtering, trends, and MinMaxLTTB.

The tradeoffs of precomputing a pyramid of extrema and statistics are worthwhile as long as visual analysis is central to working with biomedical time series.

## Availability of source code and requirements

- Project name: timeseries-zarr
- Project home page: https://github.com/Pennsieve/timeseries-zarr-py, which holds the specification and the Python reference reader and writer
- TypeScript reader: https://github.com/Pennsieve/timeseries-zarr-reader
- Operating systems: platform independent
- Programming languages: Python, TypeScript
- Other requirements: Python 3.12 or higher, zarr-python 3.2.1 or higher within version 3, NumPy 2 or higher, and numcodecs 0.16 or higher
- License: Apache-2.0
- Benchmark: https://github.com/Pennsieve/timeseries-zarr-benchmark, Python 3.12, Apache-2.0

## Data availability

The benchmarks used synthetic recordings. The benchmark code generates each one from a fixed random seed.

## Declarations

## List of abbreviations

AWS: Amazon Web Services
BIDS: Brain Imaging Data Structure
CPU: central processing unit
EDF: European Data Format
EEG: electroencephalography
EMG: electromyography
FAIR: findable, accessible, interoperable, and reusable
HDF5: Hierarchical Data Format version 5
HTTP: Hypertext Transfer Protocol
JSON: JavaScript Object Notation
LTTB: largest triangle three buckets
MEF: Multiscale Electrophysiology Format
NWB: Neurodata Without Borders
OME-NGFF: Open Microscopy Environment next-generation file format
REM: rapid eye movement
RMS: root mean square
S3: Amazon Simple Storage Service
VCF: variant call format
vCPU: virtual CPU.

## Appendix A: Format specification

Figures and tables of the appendices are numbered from S1.

### Specification details

The layout is a Zarr v3 hierarchy [27]. Each channel is a Zarr group, and its arrays are chunked along time, packed into shards so that the number of objects stays small [44], and compressed with Zstandard. Every float array declares NaN as its fill value, and a producer may omit a chunk of the samples or of a level that contains only fill. A gap that covers whole chunks is not stored. Every time in a bundle is an offset in microseconds from a single recording-wide reference time. A recording that resumes off the sample grid after a pause may carry a `runs` table. Each row describes a span of samples one sample period apart and gives the time of its first sample. The metadata a viewer needs is in one attribute, `tsz`, on each node it describes. The root also carries a consolidated copy of the metadata of every node, which a viewer reads in its first request.

Level *k* of a continuous channel has one bin for every *s*_*k*_ consecutive samples. The specification requires *s*_*k*+1_ to be a whole multiple of *s*_*k*_, at least 2*s*_*k*_, and lets the step *s*_*k*+1_*/s*_*k*_ differ from level to level. With a constant step *s, s*_*k*_ = *s*^*k*^. Each statistic is computed over the finite samples of its bin. The statistics of a bin are computed from those of the bins below it, with the mean and variance by the pairwise rule of Chan et al. [45]. Statistics computed this way match those computed from the samples, exactly for minmax and count and within a stated tolerance for the float32 mean and var. Variance was chosen in place of the sum of squares that BigWig uses [20], because a float32 sum of squares loses precision when the samples carry a large constant offset. `minmax`, `mean`, and `var` are recorded as NaN only when a bin contains no finite sample, in which case its count is zero. A bin that contains a shorter gap maintains statistics from its finite samples and its count the number of finite samples the bin contains.

Appendix B derives the displacement bound for MinMaxLTTB computed from a level and the bounds for montaged and filtered views from the level statistics.

### Event channels

An event channel stores sorted times as microsecond offsets with optional parallel arrays for labels, values, text, the channels an event applies to, and waveforms, and declares an upper bound on event duration. At a constant step *s*, the specification recommends a level 1 bin width of 1000 ·*s*^*m*^ microseconds for its count levels, with *m* an integer, so that count levels in different bundles share bin edges.

Figure S1 draws a synthetic session of 1 h, generated from a fixed random seed. Its unit channel holds 16 units that fire as Poisson processes with a 2 ms refractory period, at base rates from 1 to 20 Hz. Eight units change their rate during trials, and three drift over the hour. Each spike stores 48 samples at 30 kHz, drawn from the template of its unit with amplitude jitter and noise. The trial channel holds 598 intervals of 1.5 to 2.5 s, each with a category and a note. The reference writer wrote both channels, the count levels of the unit channel, and `waveform_mean`. It does not write a `waveforms` array. We added the array with the storage routines of the writer’s package and consolidated the metadata again.

**Figure S1:**
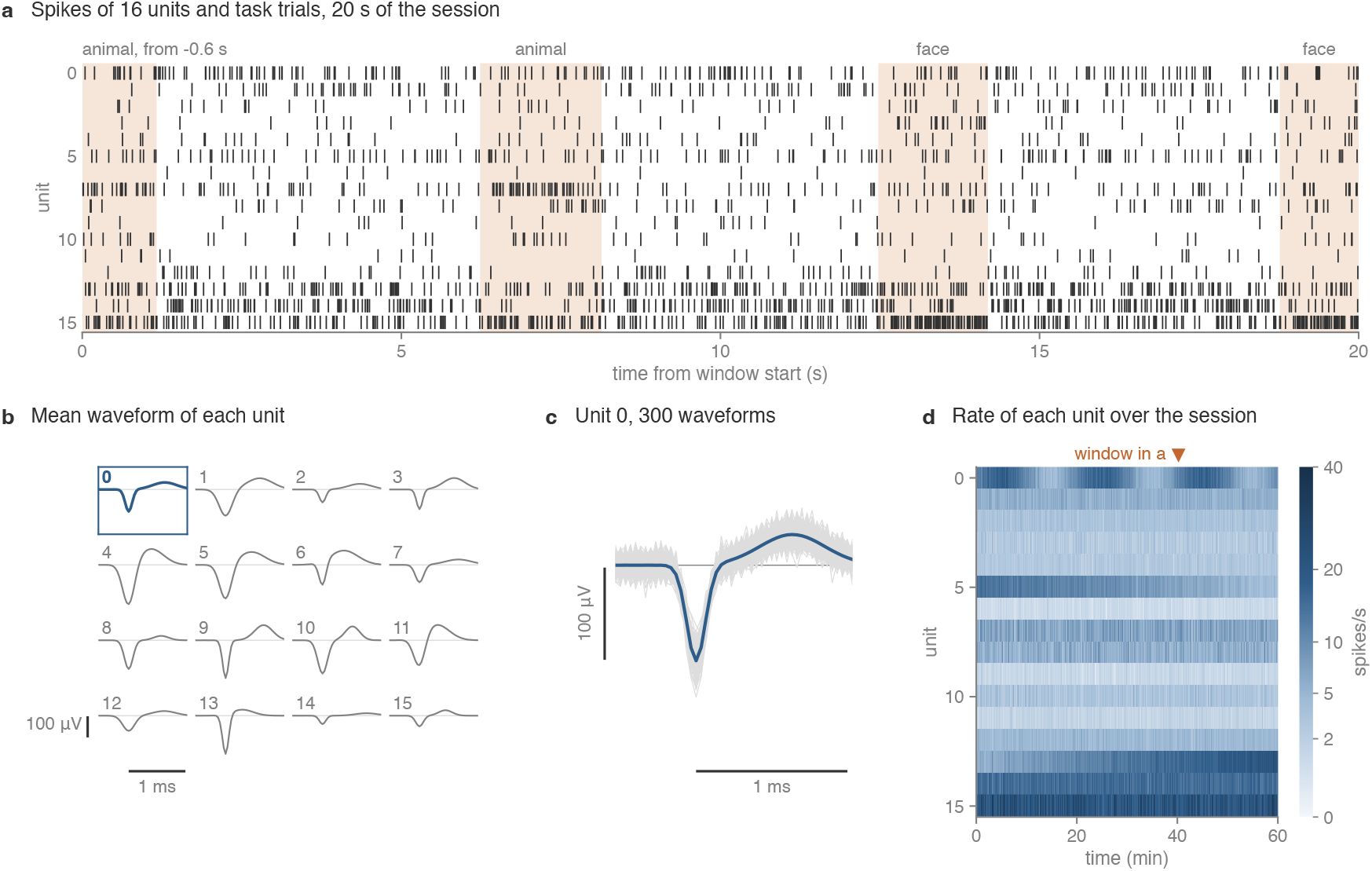
A synthetic session of 16 sorted units and 598 task trials over 1 h, stored as two event channels of one bundle. (a) A 20 s raster of the units from the event times and labels, with the trials that overlap the window shaded and named by category. The first trial began 0.6 s before the left edge and is found through the duration bound. (b) The mean waveform of each label, from `waveform_mean`. (c) 300 of the 38,439 stored waveforms of unit 0, with its mean. Scale bars in (b) and (c) mark 1 ms and 100 *µ*V. (d) The rate of each unit over the hour from count level 4, whose 3,516 bins of 1.024 s hold 225 kB before compression. Each column sums 8 bins, and the color scale is the square root of the rate. The event times and labels of the hour hold 3.8 MB before compression. The marker gives the window of (a).

## Appendix B: Error bounds on views from the level statistics

### Displacement of MinMaxLTTB computed from a level

Let a window of width *W* be drawn on *n* pixel columns, each of width *W/n*. MinMaxLTTB computed from a level uses the coarsest level with at least two bins per pixel column. Its bin width *b* satisfies *b*≤*W/*(2*n*). At a step of four, the next coarser level has bins of width 4*b > W/*(2*n*), so *b > W/*(8*n*) and each column contains fewer than eight bins. Each bin contributes its minimum and maximum, so the preselection has between four and sixteen points per column. A level stores each extremum with its bin and not with a time, so a viewer can place it at the center of the bin. The extremum lies inside the bin, so the displacement is at most *b/*2 ≤*W/*(4*n*), a quarter of a pixel column.

The minimum and maximum of a bin have the same time, the center of the bin, so their order within the bin does not affect the view. Each LTTB bucket consists of whole bins and selects one point. Because the displacement changes the triangle areas that LTTB compares, the point selected for a bucket can differ from that of exact MinMaxLTTB when two candidate areas differ by less than that change.

### Montaged and filtered views from the level statistics

Here we discuss views that can be computed exactly from stored per-bin statistics, views that can only be bounded, and how the error of the remaining views arises. We show that a montage computed from stored bin means coincides with the bin mean of the montaged signal (Theorem 1), that the minmax envelope of a montage is not determined by per-channel bin statistics and admits a conservative bound (Proposition 2). We quantify the width of that bound under a Gaussian model of the within-bin fluctuations (Theorem 3). For filtering, we decompose their error into a **redesign term** (determined by the filters alone) and an **aliasing term** (determined by the spectrum of the signal) (see Theorem 4 and its Corollary), and we give the gain of the bin mean at each frequency, which enters the aliasing term (Theorem 5). We assume the standard results in signal processing.

#### Notations

A *channel* is a sequence *x* : ℤ →R whose sample *x*[*n*] is taken at time *t*_0_ + *n/f*_*s*_, where *f*_*s*_ *>* 0 is the sampling rate in hertz and *t*_0_ is the offset. Channels lie on a *common grid* if they have the same *f*_*s*_ and the same *t*_0_. Level *k*≥ 1 has bin width *M* = 4^*k*^ samples, rate *f*_*s*_*/M* and Nyquist frequency *f*_*s*_*/*(2*M*); its bin *i* ∈ℤ is the index set *I*_*i*_ = {*M i*, …, *Mi* + *M*− 1.} The *bin mean* of level *k* is the map *D*_*M*_ defined by

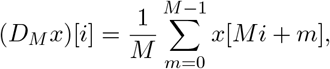

and the minmax *envelope* of *x* in bin *i* is the pair 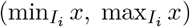. Samples are finite unless stated otherwise; non-finite samples (gaps, NaNs) are treated after Theorem 1.

#### Montaging

##### Definitions.

Let *x*_1_, …, *x*_*J*_ be channels on a common grid and *c* ∈ ℝ^*J*^ a fixed vector. The *montage* with coefficients *c* is the channel 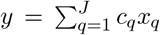. Bipolar (*c*_*a*_ = 1, *c*_*b*_ = −1, all other *c*_*q*_ = 0), common-average (*c*_*q*_ = *δ*_*qa*_ − 1*/J* for a reference channel *a*), linked-ear and Laplacian references are montages. For an interval [*u, v*] and *c* ∈ ℝ, let *c* [*u, v*] denote [*cu, cv*] if *c* ≥ 0 and [*cv, cu*] if *c <* 0. The *interval band* of *y* in bin *i* is the Minkowski sum

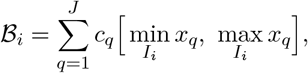

which for a bipolar pair *y* = *a* −*b* is [min *a*−max *b*, max *a*−min *b*]. A *bin statistic* is a function *S* of the vector *v* ∈ℝ^*M*^ of the samples of one bin. It is *permutation-invariant* if *S*(*v* ◦*π*) = *S*(*v*) for every *v* and every permutation *π* of {0, …, *M*− 1}. The stored minimum, maximum, mean, variance and count are permutation-invariant, within a bin.

##### Theorem 1

(Linear views are exact). *Let L be a linear map on real sequences, applied identically to every channel: for example D*_*M*_, *a linear filter, or a composition of these. Then every montage y* = ∑_*q*_ *c*_*q*_*x*_*q*_ *satisfies L*(*y*) = ∑_*q*_ *c*_*q*_ *L*(*x*_*q*_).

*Proof*. Linearity of *L*, applied to a finite sum. □

The content of the theorem lies in its hypotheses, which a viewer must check. (a) *Common grid*. Channels with different rates have no common grid. If two offsets differ by *s/f*_*s*_ with *s ∈*ℤ, re-indexing puts the raw samples on a common grid, but the bins of level *k* of the two channels then cover the same times only if *M* divides *s*; otherwise *D*_*M*_ is not applied identically and the identity fails. (b) *Filter state*. Filters in the sense of Section, which include recursive filters run in steady state, are linear, and so is recursive filtering started from a zero state at a fixed index. Filtering from a fixed nonzero initial state is affine, and the identity then holds only if the initial state used for *y* is ∑_*q*_ *c*_*q*_*s*_*q*_, where *s*_*q*_ is the one used for *x*_*q*_, as it is when every state is computed from preceding data. (c) *Gaps*. Represent a non-finite sample by NaN, absorbing under addition and under multiplication by a nonzero scalar, and let the montage sum only over *q* with *c*_*q*_ ≠0. For the NaN-propagating bin mean, which is NaN on every bin that contains a NaN, both sides of the identity are NaN exactly when some *x*_*q*_ with *c*_*q*_ ≠ 0 has a gap in *I*_*i*_, and they are equal otherwise. A mean that skips non-finite samples breaks the identity: with *M* = 2, *a* = (1, 3) and *b* = (NaN, 1), the skipping mean of *a b* = (NaN, 2) is 2, whereas the difference of the skipping means is 2− 1 = 1. The stored fold propagates NaN, so a stored mean level equals *D*_*M*_ applied to raw in this sense.

##### Proposition 2

(The montaged `minmax` envelope is not recoverable). *Let bins have M*≥ 2 *samples. (i) There are bins a, b and b*′ *such that S*(*b*) = *S*(*b*′) *for every permutation-invariant statistic S, but* max(*a − b*) ≠ max(*a − b*′) *and* min(*a − b*) ≠ min(*a − b*′); *hence no function of per-channel bin statistics gives the minimum or maximum of a montage. (ii) Every montage satisfies y*[*m*] ∈ℬ_*i*_ *for all m ∈ I*_*i*_, *and for every montage there are permutations of each channel’s samples within the bin, which leave every permutation-invariant statistic unchanged, after which y attains both endpoints of ℬ*_*i*_. *Consequently, any interval computed from permutation-invariant per-channel statistics that contains y on I*_*i*_ *for all inputs contains ℬ*_*i*_; *for a bipolar pair*, [min *a* −max *b*, max *a*− min *b*] *is the tightest such interval*.

*Proof*. (i) Take *a* = *b* = (0, 10) and *b*′ = (10, 0), and for *M >* 2 append *M*− 2 samples equal to 5 to each. Then *b*′ is a permutation of *b*, while *a* −*b* is identically 0 and *a* −*b*′ takes the values −10 and 10. (ii) For *m* ∈ *I*_*i*_, 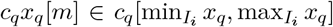 for every *q*; summing over *q* gives *y*[*m*] ∈ℬ_*i*_. Choose distinct indices *m*_*−*_, *m*_+_ *I*_*i*_, which exist because *M ≥*2. Permute the samples of each *x*_*q*_ within the bin so that *x*_*q*_[*m*_*−*_] is its minimum and *x*_*q*_[*m*_+_] its maximum if *c*_*q≥*_ 0, and the reverse if *c*_*q*_ *<* 0. Then *y*[*m*_*−*_] and *y*[*m*_+_] are the lower and upper endpoints of ℬ_*i*_. An interval computed from the statistics is the same before and after the permutation, so if it is to contain *y* in both cases it contains both endpoints, and therefore ℬ_*i*_. □

##### Theorem 3

(Width of the interval band). *Let a bin hold M* ≥ 2 *samples, and let the pairs* (*a*[*m*], *b*[*m*]), *m* ∈ *I*_*i*_, *be independent and identically distributed, bivariate Gaussian with arbitrary means, common variance σ*^2^ *>* 0 *and correlation ρ*_*w*_ ∈ [−1, 1). *Let W*_*ℬ*_ = (max *a*−min *a*)+(max *b*− min *b*) *be the width of the interval band of a* − *b and W* = max(*a* −*b*) − min(*a* − *b*) *the width of its minmax envelope, all extrema taken over I*_*i*_. *Then* 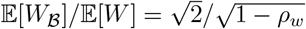.

*Proof*. Let *c*_*M*_∈(0, ∞) be the expected range of *M* independent standard Gaussian variables. The range of *M* independent *N*(*µ, s*^2^) variables with *s >* 0 is *s* times the range of standard ones, so its expectation is *c*_*M*_ *s*. The samples of *a* are independent *N*(*µ*_*a*_, *σ*^2^), and likewise for *b*; by linearity of expectation E[*W*_*B*_] = 2*c*_*M*_ *σ*, whatever the dependence between *a* and *b*. The samples of *a b* are independent *N*(*µ*− *µ*, 2*σ*^2^(1 −*ρ*)), with positive variance because *ρ*_*w*_ *<* 1, so 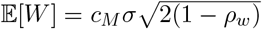. The constant *c*_*M*_ cancels in the ratio. □

#### Filtering

**Definitions**. On a grid of rate *R* (below, either the native rate *f*_*s*_ or a level rate *f*_*s*_*/M*), a frequency *f* in hertz corresponds to the normalized angular frequency *ω* = 2*πf/R*, taken modulo 2*π* in [−*π, π*). For *x* ∈ *l*^2^(ℤ) the discrete-time Fourier transform is *X*(*ω*) = (ℱ*x*)(*ω*) = ∑_*n*_ *x*[*n*]*e*^*−jωn*^, and 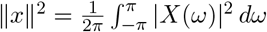. A *filter* is a linear, time-invariant map *G* on *l*^2^(ℤ) given by ℱ[*Gx*](*ω*) = *G*(*ω*)*X*(*ω*), where the *frequency response G*(*ω*) is a bounded, measurable, 2*π*-periodic function. We write *G*(*f*) for the response at the physical frequency *f*.

The bin mean factors as *D*_*M*_ = ↓_*M*_ ◦ (*h*_*B*_ * ·), where the *boxcar h*_*B*_ is 1*/M* for −*M < n* ≤ 0 and 0 otherwise, and (↓_*M*_ *v*)[*i*] = *v*[*Mi*] is *downsampling*. The boxcar has frequency response

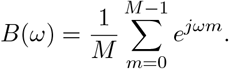

Let *H* be a filter at the native rate *f*_*s*_, with frequency variable *ω*, and *H*_*l*_ a filter at the level rate *f*_*s*_*/M*, with frequency variable Ω; one physical frequency has Ω = *Mω*. For Ω∈ [− *π, π*) the *fold frequencies ω*_*r*_ = (Ω + 2*πr*)*/M, r* = 0, …, *M −*1, are the native frequencies that downsampling by *M* maps onto Ω. Let *f*_max_ be the highest critical (transition) frequency of the filter specification. The *transition ratio* at level *k* is

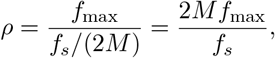

and a design at the level rate requires *ρ <* 1. We compare *D*_*M*_ (*Hx*), the native-rate filter followed by the bin mean, with *H*_*l*_(*D*_*M*_ *x*), the level-rate filter applied to stored means. Both end at the level rate, so the comparison isolates the order of operations.

##### Theorem 4

(Error decomposition). *Let x* ∈ *l*^2^(ℤ), *and let H and H*_*l*_ *be filters at the native and the level rate. For almost every* Ω ∈ [−*π, π*),

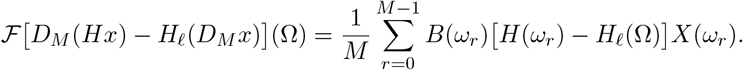

*The term r* = 0 *is the* redesign term *and the terms r* ≥ 1 *are the* aliasing terms; *e*_red_ *and e*_alias_ *denote the corresponding parts of the error*.

*Proof*. For *v l*^2^(ℤ) write *V* = ℱ*v*. By the definition of the fold frequencies, *Mω* = Ω + 2*πr*, ∈ ℱ _*r*_ so *Mω*_*r*_ ≡ Ω (mod 2*π*) for every *r*, with equality for *r* = 0. Since 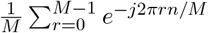 is 1 if *M* divides *n* and 0 otherwise,

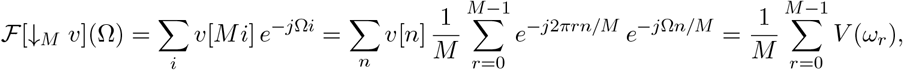

where the series converge in *L*^2^, so the identity holds for almost every Ω. By the convolution theorem, *v* = *h*_*B*_ * *Hx* has transform *BHX* and *v* = *h*_*B*_ * *x* has transform *BX*; applying the identity to each gives

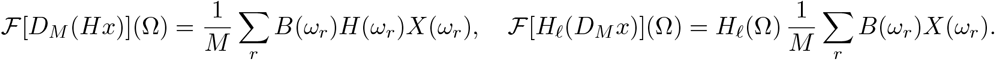

Subtract and group the terms by *r*. □

##### Corollary.

Let Π_in_ and Π_out_ be the orthogonal projections of *l*^2^(ℤ) onto the frequencies *ω < π/M* and *π/M≤* |*ω*| *≤π*, and let *C*_*H*_ be the essential supremum of |*H*(*ω*_*r*_) −*H*_*l*_(Ω)| over Ω∈ [−*π, π*) and 1 *≤ r ≤ M*− 1, so *C*_*H*_ *≤* sup |*H*| + sup |*H*_*l*_|. (i) If *H*_*l*_(Ω) = *H*(Ω*/M*) for Ω ∈ [−*π, π*), that is, if *H*_*l*_ matches the native response, then *e*_red_ = 0. (ii) ∥*e*_red_ ∥*M* ^*−*1*/*2^ sup_|*ω*|*<π/M*_ |*H*(*ω*) −*H*_*l*_(*Mω*)| ∥Π_in_(*h*_*B*_ *x*) ∥. (iii) *e*_alias_ *≤ C*_*H*_ *∥*Π_out_(*h*_*B*_ *x*) ∥.

*Proof*. (i) is immediate. For (ii), write Δ(*ω*) = *H*(*ω*) −*H*_*l*_(*Mω*). Parseval’s identity applied to the term *r* = 0 and the substitution *ω* = Ω*/M, d*Ω = *M dω*, give

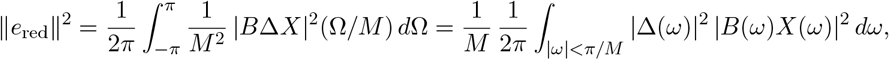

and bounding |Δ| by its supremum leaves ∥Π_in_(*h*_*B*_ *\*x*) ∥ ^2^, since *h*_*B*_ *\*x* has transform *BX*. The factor *M* ^*−*1*/*2^∥Π_in_(*h*_*B*_*\*x*)∥ equals ∥ ↓_*M*_ Π_in_(*h*_*B \**_ *x*) ∥, by the downsampling identity in the proof of Theorem 4 with only the term *r* = 0 nonzero; it is the norm of the part of the stored mean level *D*_*M*_ *x* that comes from frequencies below the Nyquist frequency of the level, so relative to that part the redesign error is at most the largest in-band mismatch of the two responses. For (iii), bound each bracket by *C*_*H*_ and apply the Cauchy–Schwarz inequality to the *M* − 1 terms:

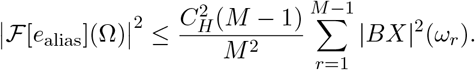

For each *r*, the substitution *ω* = *ω*_*r*_, *d*Ω = *M dω*, maps [−*π, π*) onto [(2*r −*1)*π/M*, (2*r* + 1)*π/M*). These intervals tile [*π/M*, 2*π− π/M*), which modulo 2*π* is {*π/M ≤*|*ω*|≤ π} up to a null set, and *BX* is 2*π*-periodic. Integrating over Ω therefore gives

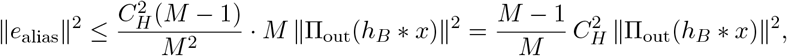

and (*M* −1)*/M* ≤1. □

The redesign error comes from the two filters disagreeing on frequencies the level can represent, so it depends only on the filters and vanishes when they match. The aliasing error comes from signal content above the Nyquist frequency of the level, folded into its band when the means were stored, so its size is set by the spectrum of the signal and, in general, only finer data can reduce it.

#### Quantities to estimate

The Corollary reduces the error of a filtered view served from level *k* to two quantities. The in-band mismatch sup_|*ω*|*<π/M*_ |*H*(*ω*) −*H*_*l*_(*Mω*)| depends only on the two filters and the level, so a viewer can compute it before reading any data. The out-of-band energy ∥Π_out_(*h*_*B*_* *x*) ∥ depends on the signal, and a bundle stores no spectral description from which to compute it.

#### The size of the boxcar factor

The aliasing terms see the input only through *B*(*ω*_*r*_)*X*(*ω*_*r*_) at the fold frequencies *ω*_*r*_, *r* ≥1, which lie above the Nyquist frequency of the level. How strongly the bin mean suppresses each frequency therefore decides how much signal content reaches them. The following result gives that suppression for a single tone.

##### Theorem 5

(Attenuation and nulls of the bin mean). *Let* 0 *< f < f, let θ* ∈ ℝ *and let x*[*n*] = cos(2*πfn/f*_*s*_+*θ*). *Then* (*D*_*M*_ *x*)[*i*] = *B*(2*πf/f*_*s*_) cos (2*πf M i/f*_*s*_+*θ*+arg *B*(2*πf/f*_*s*_) *(any argument if B* = 0*), a tone sampled at the level rate, with gain*

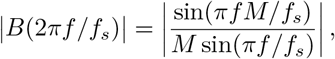

*which vanishes if and only if f is an integer multiple of the level rate f*_*s*_*/M*.

*Proof*. Let *ω* = 2*πf/f*_*s*_. For the complex tone *e*^*jωn*^, 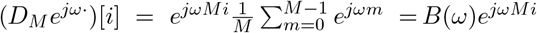; the real tone is Re (*e*^*jθ*^*e*^*jωn*^) and *D*_*M*_ is real-linear. Summing the geometric series, *B*(*ω*) = *e*^*jω*(*M−*1)*/*2^ sin(*M ω/*2)*/*(*M* sin(*ω/*2)), where sin(*ω/*2) ≠0 because 0 *< f < f*_*s*_; the numerator vanishes exactly when *f M/f*_*s*_ is an integer. □

At the level rate the output tone appears at *f* modulo *f*_*s*_*/M*, which is aliasing when *f > f*_*s*_*/*(2*M*). The closed form also lets a viewer compensate the droop of a mean level near its Nyquist frequency.

## Appendix C: Benchmarks

This appendix describes the benchmark methodology and reports the results for each recording and store.

### Recordings and hardware

The benchmark generates and reads four synthetic recordings from fixed random seeds (Table S1). R1 is one channel of ambulatory EEG at 256 Hz for 24 h. R2 is 32 channels of scalp EEG at 512 Hz for 24 h. R3 is 200 channels of intracranial EEG at 2 kHz for 24 h, the recording of Table 1. R4 is 384 channels at 30 kHz for 10 min, the channel count and rate of a Neuropixels probe. Each channel is Gaussian noise with a 1*/f* spectrum and a bump at 8 to 12 Hz, or at 300 to 3000 Hz in R4. Channel 0 of each recording has three planted gaps of NaN samples, the longest 10 min in R1 to R3 and 1 min in R4. Samples are quantized to a 0.125 *µ*V grid. The 16-bit EDF copy and the float32 copies store the same values. The format benchmark reads the first hour of R1 to R3 and all of R4, and the resampling benchmark reads every recording at full length.

**Table S1:** The synthetic recordings. The float32 size column is the size of every sample stored as float32, in decimal units. The levels column is the number of levels that the reference writer creates by default.

| id | modality | channels | rate | duration | samples per channel | float32 size | levels |
|---|---|---|---|---|---|---|---|
| R1 | ambulatory EEG | 1 | 256 Hz | 24 h | 22,118,400 | 88.5 MB | 7 |
| R2 | scalp EEG | 32 | 512 Hz | 24 h | 44,236,800 | 5.66 GB | 7 |
| R3 | intracranial EEG | 200 | 2 kHz | 24 h | 172,800,000 | 138 GB | 8 |
| R4 | Neuropixels-like | 384 | 30 kHz | 10 min | 18,000,000 | 27.6 GB | 7 |

Both benchmarks ran in one AWS region on c6i.8xlarge instances. Each instance has 32 vCPU, 64 GiB of memory, and a 12.5 Gbps network. One instance stored a local copy of each recording and ran the format benchmark. A second instance served every file over HTTP/1.1 from nginx. Both instances used gp3 volumes provisioned at 1000 MiB/s. S3 was in the same region and was read with signed requests through a gateway endpoint. The page cache was dropped on both instances before each process of the format benchmark, such that every format and store started from a cold cache and was timed under the same conditions.

### Format benchmark

Every recording read by the format benchmark was written in each format, except R4, which was not written as EDF because a record at that rate and channel count would exceed the 61,440 bytes that the EDF specification recommends. Each record of an EDF file contains every channel, and the files are uncompressed. Each NWB file stores samples in one electrical series, written once in chunks of about 1 MB along time across every channel and once in chunks of 8192 samples of one channel. Each NWB file on Zarr was a Zarr v3 store sharded like TS-Zarr, with as many whole chunks along time in each shard as fit in 16 MiB, so that each shard was one object. TS-Zarr stored every statistic in inner chunks of 8192 elements and shards of 16 MiB, and the writer stopped adding levels when the next would have fewer than 1024 bins. Each recording also had a second TS-Zarr copy of the samples alone, for a like-for-like comparison of storage against EDF and NWB. Every chunked array was compressed with Zstandard at compression level 3. The benchmark included a range reader for EDF, since no open-source EDF library read a window by byte range.

The format benchmark read windows of 1 s, 10 s, 1 min, and 10 min at random positions that were shared across formats and stores. Each combination of recording, store, format, read kind, and window ran in a fresh process that opened the file before the timer started. The timer ran from the read call to the decoded array in memory. Each combination made 100 reads, or 20 at 10 min, and stopped early after the read that brought its total read time to at least 60 s. The benchmark recorded the duration, the bytes transferred, and the number of requests of each read. Every format was read through the same filesystem layer, which allowed at most 32 concurrent requests. A 1-byte object read provided a baseline latency for each store. Reads of more than 8 GB were not measured, which excluded the all-channel read of R4 at 10 min.

The time to open a file was recorded independently of a read. On S3 the open took 5.5 to 6.1 s and 175 requests for NWB on HDF5, 1.0 to 1.5 s and 23 requests for NWB on Zarr, and two requests for TS-Zarr, whose root carries a consolidated copy of the metadata. A reader parses that metadata when it opens a recording. The open time therefore grows with the number of channels, from 0.83 s for one channel to 5.1 s for the 384 channels of R4.

**Table S2:**
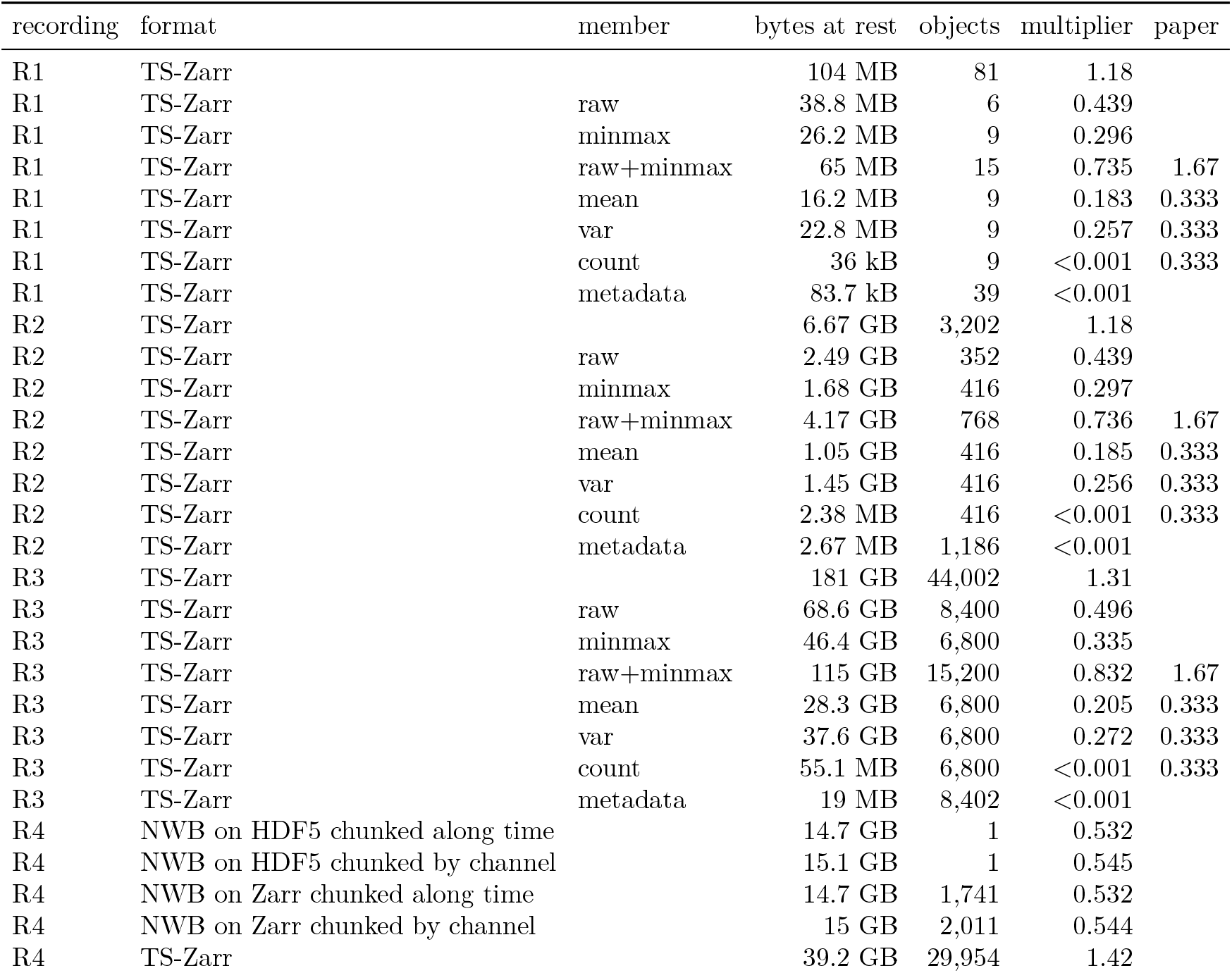
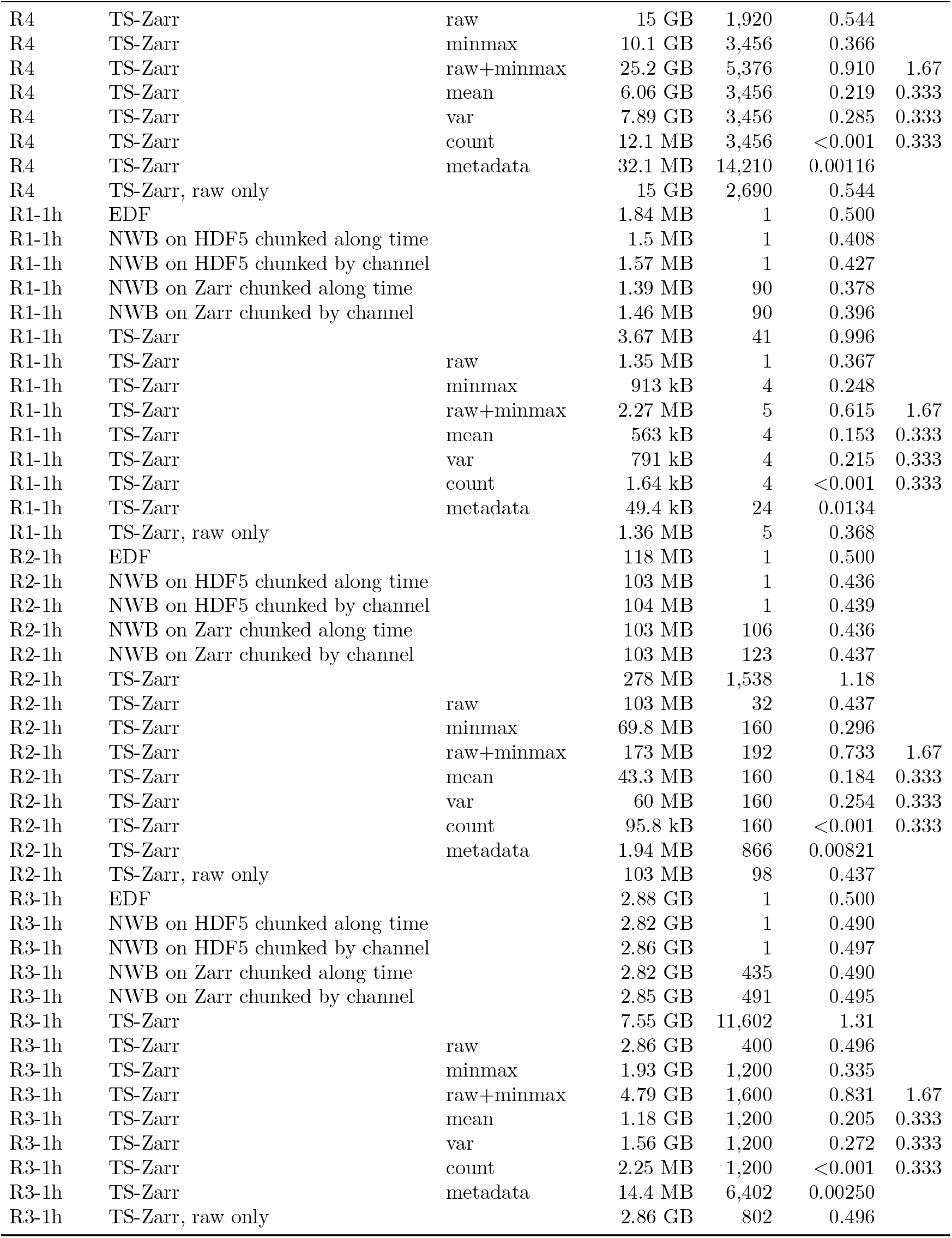
Bytes at rest, objects, and multiplier per recording and format. The multiplier is the bytes divided by the float32 bytes of the recording. The paper column gives the multiplier of the values in the main text: the samples and minmax together, and each further statistic. The raw-only row is TS-Zarr with the samples alone. R1 to R3 at full length are written as TS-Zarr only.

| recording | format | member | bytes at rest | objects | multiplier | paper |
|---|---|---|---|---|---|---|
| R1 | TS-Zarr |  | 104 MB | 81 | 1.18 |  |
| R1 | TS-Zarr | raw | 38.8 MB | 6 | 0.439 |  |
| R1 | TS-Zarr | minmax | 26.2 MB | 9 | 0.296 |  |
| R1 | TS-Zarr | raw+minmax | 65 MB | 15 | 0.735 | 1.67 |
| R1 | TS-Zarr | mean | 16.2 MB | 9 | 0.183 | 0.333 |
| R1 | TS-Zarr | var | 22.8 MB | 9 | 0.257 | 0.333 |
| R1 | TS-Zarr | count | 36 kB | 9 | <0.001 | 0.333 |
| R1 | TS-Zarr | metadata | 83.7 kB | 39 | <0.001 |  |
| R2 | TS-Zarr |  | 6.67 GB | 3,202 | 1.18 |  |
| R2 | TS-Zarr | raw | 2.49 GB | 352 | 0.439 |  |
| R2 | TS-Zarr | minmax | 1.68 GB | 416 | 0.297 |  |
| R2 | TS-Zarr | raw+minmax | 4.17 GB | 768 | 0.736 | 1.67 |
| R2 | TS-Zarr | mean | 1.05 GB | 416 | 0.185 | 0.333 |
| R2 | TS-Zarr | var | 1.45 GB | 416 | 0.256 | 0.333 |
| R2 | TS-Zarr | count | 2.38 MB | 416 | <0.001 | 0.333 |
| R2 | TS-Zarr | metadata | 2.67 MB | 1,186 | <0.001 |  |
| R3 | TS-Zarr |  | 181 GB | 44,002 | 1.31 |  |
| R3 | TS-Zarr | raw | 68.6 GB | 8,400 | 0.496 |  |
| R3 | TS-Zarr | minmax | 46.4 GB | 6,800 | 0.335 |  |
| R3 | TS-Zarr | raw+minmax | 115 GB | 15,200 | 0.832 | 1.67 |
| R3 | TS-Zarr | mean | 28.3 GB | 6,800 | 0.205 | 0.333 |
| R3 | TS-Zarr | var | 37.6 GB | 6,800 | 0.272 | 0.333 |
| R3 | TS-Zarr | count | 55.1 MB | 6,800 | <0.001 | 0.333 |
| R3 | TS-Zarr | metadata | 19 MB | 8,402 | <0.001 |  |
| R4 | NWB on HDF5 chunked along time |  | 14.7 GB | 1 | 0.532 |  |
| R4 | NWB on HDF5 chunked by channel |  | 15.1 GB | 1 | 0.545 |  |
| R4 | NWB on Zarr chunked along time |  | 14.7 GB | 1,741 | 0.532 |  |
| R4 | NWB on Zarr chunked by channel |  | 15 GB | 2,011 | 0.544 |  |
| R4 | TS-Zarr |  | 39.2 GB | 29,954 | 1.42 |  |
| R4 | TS-Zarr | raw | 15 GB | 1,920 | 0.544 |  |
| R4 | TS-Zarr | minmax | 10.1 GB | 3,456 | 0.366 |  |
| R4 | TS-Zarr | raw+minmax | 25.2 GB | 5,376 | 0.910 | 1.67 |
| R4 | TS-Zarr | mean | 6.06 GB | 3,456 | 0.219 | 0.333 |
| R4 | TS-Zarr | var | 7.89 GB | 3,456 | 0.285 | 0.333 |
| R4 | TS-Zarr | count | 12.1 MB | 3,456 | <0.001 | 0.333 |
| R4 | TS-Zarr | metadata | 32.1 MB | 14,210 | 0.00116 |  |
| R4 | TS-Zarr, raw only |  | 15 GB | 2,690 | 0.544 |  |
| R1-1h | EDF |  | 1.84 MB | 1 | 0.500 |  |
| R1-1h | NWB on HDF5 chunked along time |  | 1.5 MB | 1 | 0.408 |  |
| R1-1h | NWB on HDF5 chunked by channel |  | 1.57 MB | 1 | 0.427 |  |
| R1-1h | NWB on Zarr chunked along time |  | 1.39 MB | 90 | 0.378 |  |
| R1-1h | NWB on Zarr chunked by channel |  | 1.46 MB | 90 | 0.396 |  |
| R1-1h | TS-Zarr |  | 3.67 MB | 41 | 0.996 |  |
| R1-1h | TS-Zarr | raw | 1.35 MB | 1 | 0.367 |  |
| R1-1h | TS-Zarr | minmax | 913 kB | 4 | 0.248 |  |
| R1-1h | TS-Zarr | raw+minmax | 2.27 MB | 5 | 0.615 | 1.67 |
| R1-1h | TS-Zarr | mean | 563 kB | 4 | 0.153 | 0.333 |
| R1-1h | TS-Zarr | var | 791 kB | 4 | 0.215 | 0.333 |
| R1-1h | TS-Zarr | count | 1.64 kB | 4 | <0.001 | 0.333 |
| R1-1h | TS-Zarr | metadata | 49.4 kB | 24 | 0.0134 |  |
| R1-1h | TS-Zarr, raw only |  | 1.36 MB | 5 | 0.368 |  |
| R2-1h | EDF |  | 118 MB | 1 | 0.500 |  |
| R2-1h | NWB on HDF5 chunked along time |  | 103 MB | 1 | 0.436 |  |
| R2-1h | NWB on HDF5 chunked by channel |  | 104 MB | 1 | 0.439 |  |
| R2-1h | NWB on Zarr chunked along time |  | 103 MB | 106 | 0.436 |  |
| R2-1h | NWB on Zarr chunked by channel |  | 103 MB | 123 | 0.437 |  |
| R2-1h | TS-Zarr |  | 278 MB | 1,538 | 1.18 |  |
| R2-1h | TS-Zarr | raw | 103 MB | 32 | 0.437 |  |
| R2-1h | TS-Zarr | minmax | 69.8 MB | 160 | 0.296 |  |
| R2-1h | TS-Zarr | raw+minmax | 173 MB | 192 | 0.733 | 1.67 |
| R2-1h | TS-Zarr | mean | 43.3 MB | 160 | 0.184 | 0.333 |
| R2-1h | TS-Zarr | var | 60 MB | 160 | 0.254 | 0.333 |
| R2-1h | TS-Zarr | count | 95.8 kB | 160 | <0.001 | 0.333 |
| R2-1h | TS-Zarr | metadata | 1.94 MB | 866 | 0.00821 |  |
| R2-1h | TS-Zarr, raw only |  | 103 MB | 98 | 0.437 |  |
| R3-1h | EDF |  | 2.88 GB | 1 | 0.500 |  |
| R3-1h | NWB on HDF5 chunked along time |  | 2.82 GB | 1 | 0.490 |  |
| R3-1h | NWB on HDF5 chunked by channel |  | 2.86 GB | 1 | 0.497 |  |
| R3-1h | NWB on Zarr chunked along time |  | 2.82 GB | 435 | 0.490 |  |
| R3-1h | NWB on Zarr chunked by channel |  | 2.85 GB | 491 | 0.495 |  |
| R3-1h | TS-Zarr |  | 7.55 GB | 11,602 | 1.31 |  |
| R3-1h | TS-Zarr | raw | 2.86 GB | 400 | 0.496 |  |
| R3-1h | TS-Zarr | minmax | 1.93 GB | 1,200 | 0.335 |  |
| R3-1h | TS-Zarr | raw+minmax | 4.79 GB | 1,600 | 0.831 | 1.67 |
| R3-1h | TS-Zarr | mean | 1.18 GB | 1,200 | 0.205 | 0.333 |
| R3-1h | TS-Zarr | var | 1.56 GB | 1,200 | 0.272 | 0.333 |
| R3-1h | TS-Zarr | count | 2.25 MB | 1,200 | <0.001 | 0.333 |
| R3-1h | TS-Zarr | metadata | 14.4 MB | 6,402 | 0.00250 |  |
| R3-1h | TS-Zarr, raw only |  | 2.86 GB | 802 | 0.496 |  |

**Figure S2:**
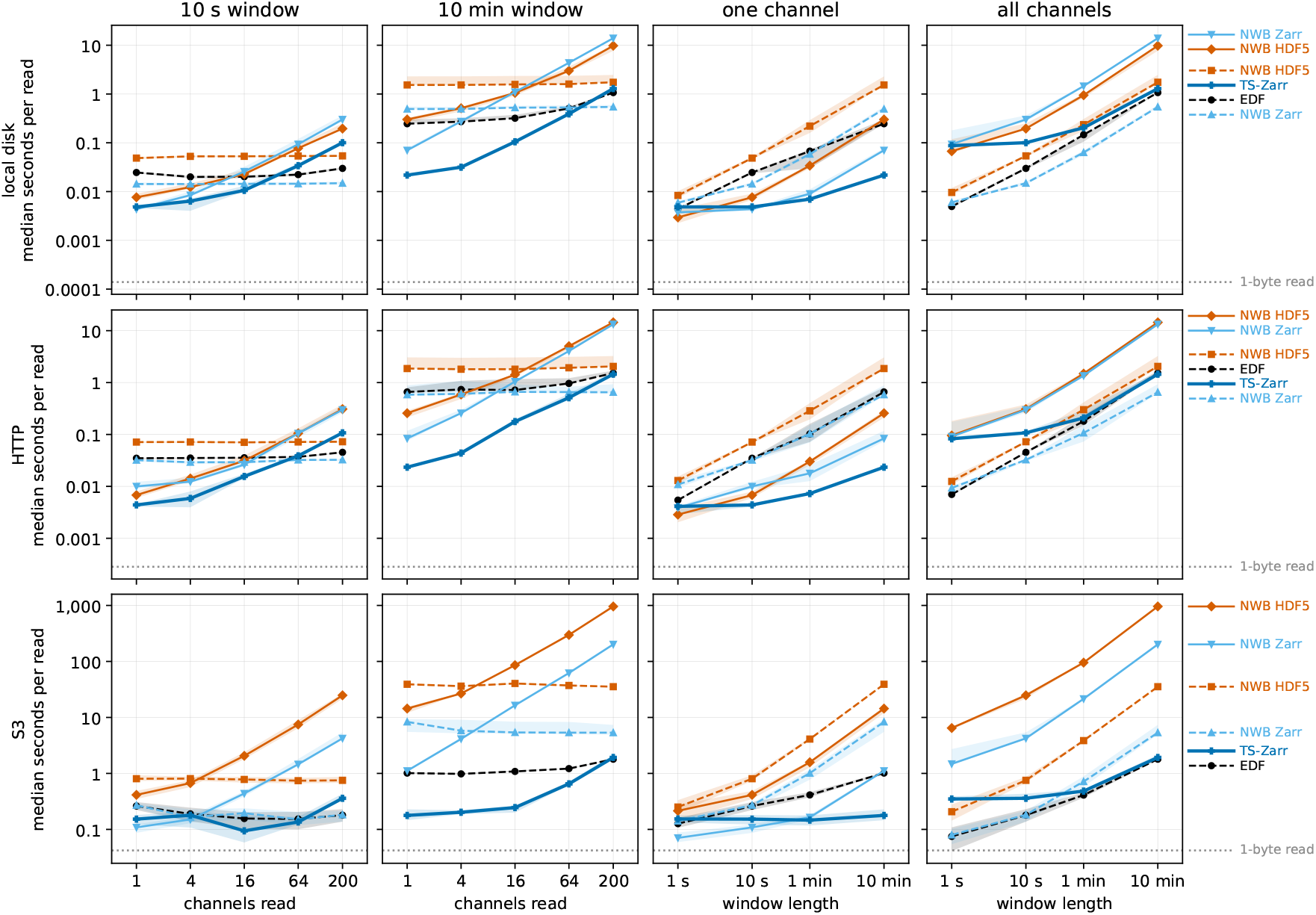
Median seconds per read of the raw samples of the first hour of R3 on local disk, behind HTTP, and on S3, on log axes, one row per store. The channel panels read windows of 10 s and 10 min from a growing number of channels. The window panels read one channel and all channels at growing window lengths. Solid lines are formats that store each channel separately (TS-Zarr and the NWB files chunked by channel). Dashed lines are formats that store a span of time of every channel together (EDF and the NWB files chunked along time). A TS-Zarr read fetches the raw arrays and no level. Bands span the 25th to 75th percentile of the reads, and the dotted line is the median of a 1-byte read from the same store. The medians cover 5,873 reads on local disk, 5,844 on HTTP, and 4,535 on S3. Each combination of format, read, and window has between one and 100 reads. It stops after the read that brings its total read time to at least 60 s.

**Figure S3:**
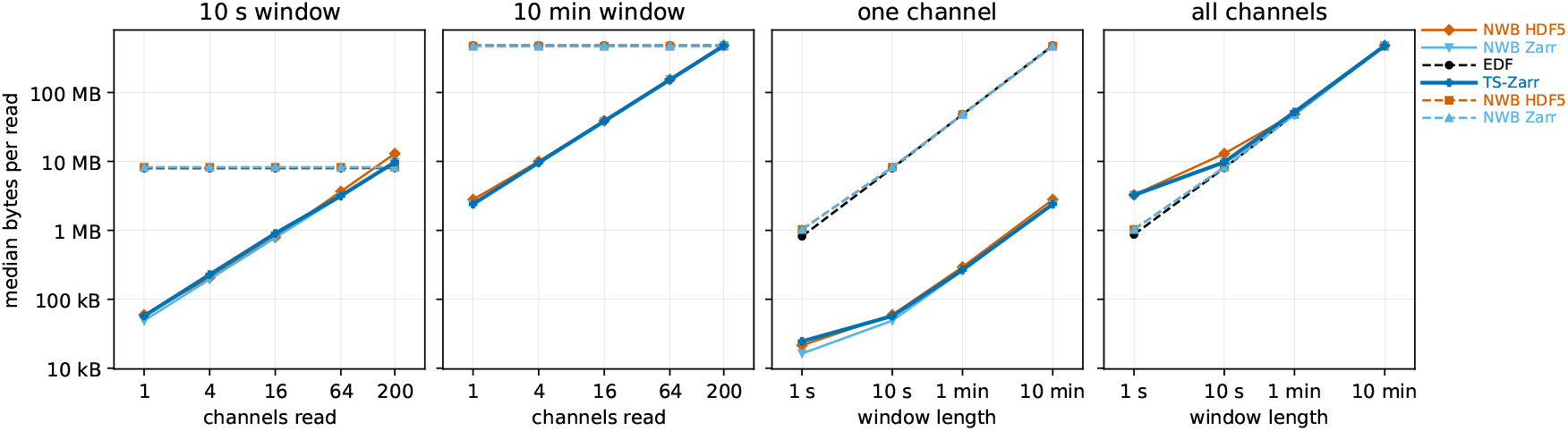
Median bytes that S3 served per read of the raw samples of the first hour of R3, on log axes. The bytes do not depend on the store. The channel panels read windows of 10 s and 10 min from a growing number of channels. The window panels read one channel and all channels at growing window lengths. The medians cover 4,535 reads. Solid lines are formats that store each channel separately (TS-Zarr and the NWB files chunked by channel). Dashed lines are formats that store a span of time of every channel together (EDF and the NWB files chunked along time). A TS-Zarr read fetches the raw arrays and no level. Each combination of format, read, and window has between one and 100 reads. It stops after the read that brings its total read time to at least 60 s.

**Figure S4:**
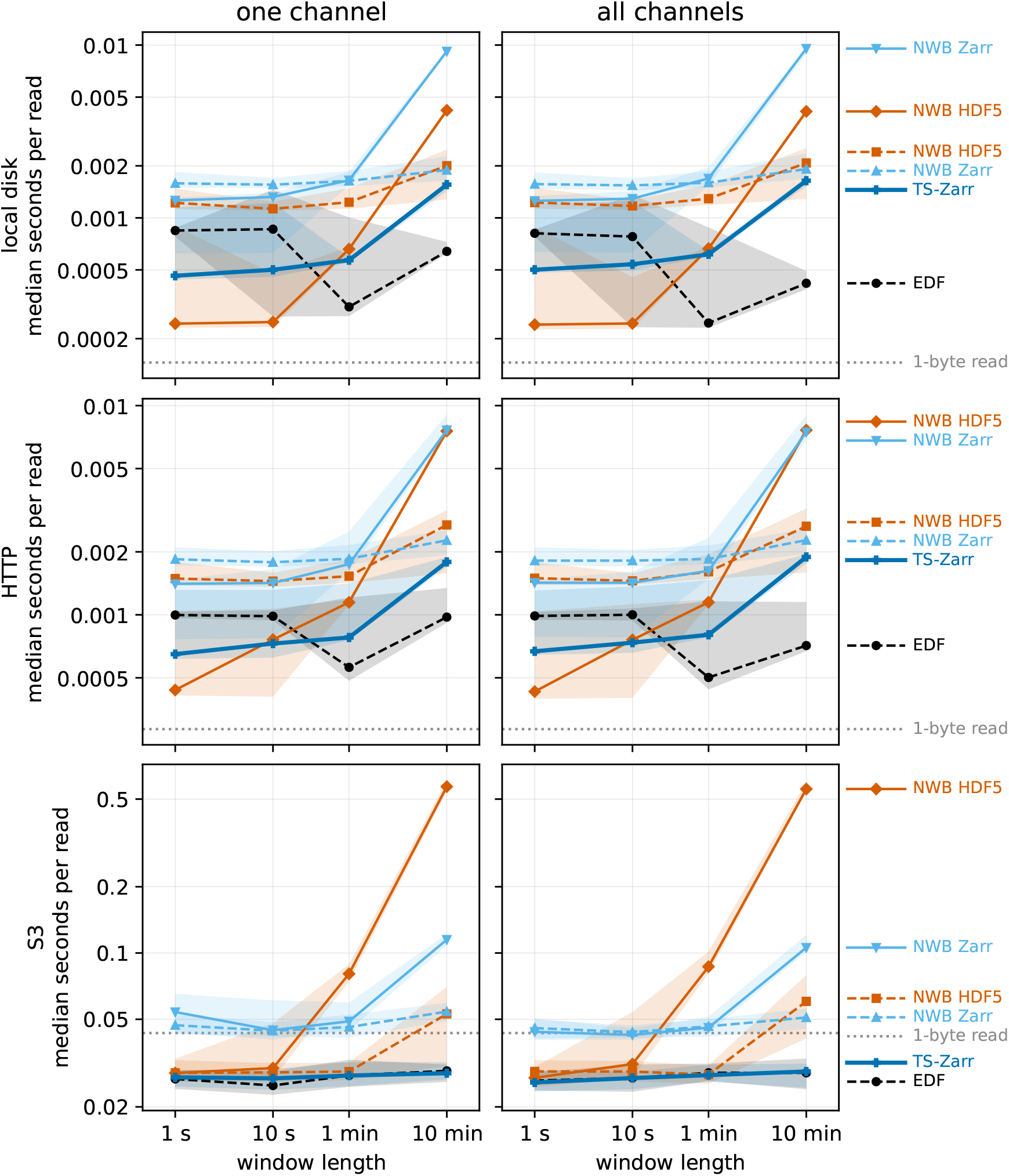
Median seconds per read of the raw samples of the first hour of R1 on local disk, behind HTTP, and on S3, on log axes. Only the window panels are drawn, since R1 has one channel. Solid lines are formats that store each channel separately (TS-Zarr and the NWB files chunked by channel). Dashed lines are formats that store a span of time of every channel together (EDF and the NWB files chunked along time). A TS-Zarr read fetches the raw arrays and no level. Bands span the 25th to 75th percentile of the reads, and the dotted line is the median of a 1-byte read from the same store. The medians cover 3,840 reads on each store. Each combination of format, read, and window has between one and 100 reads. It stops after the read that brings its total read time to at least 60 s.

**Figure S5:**
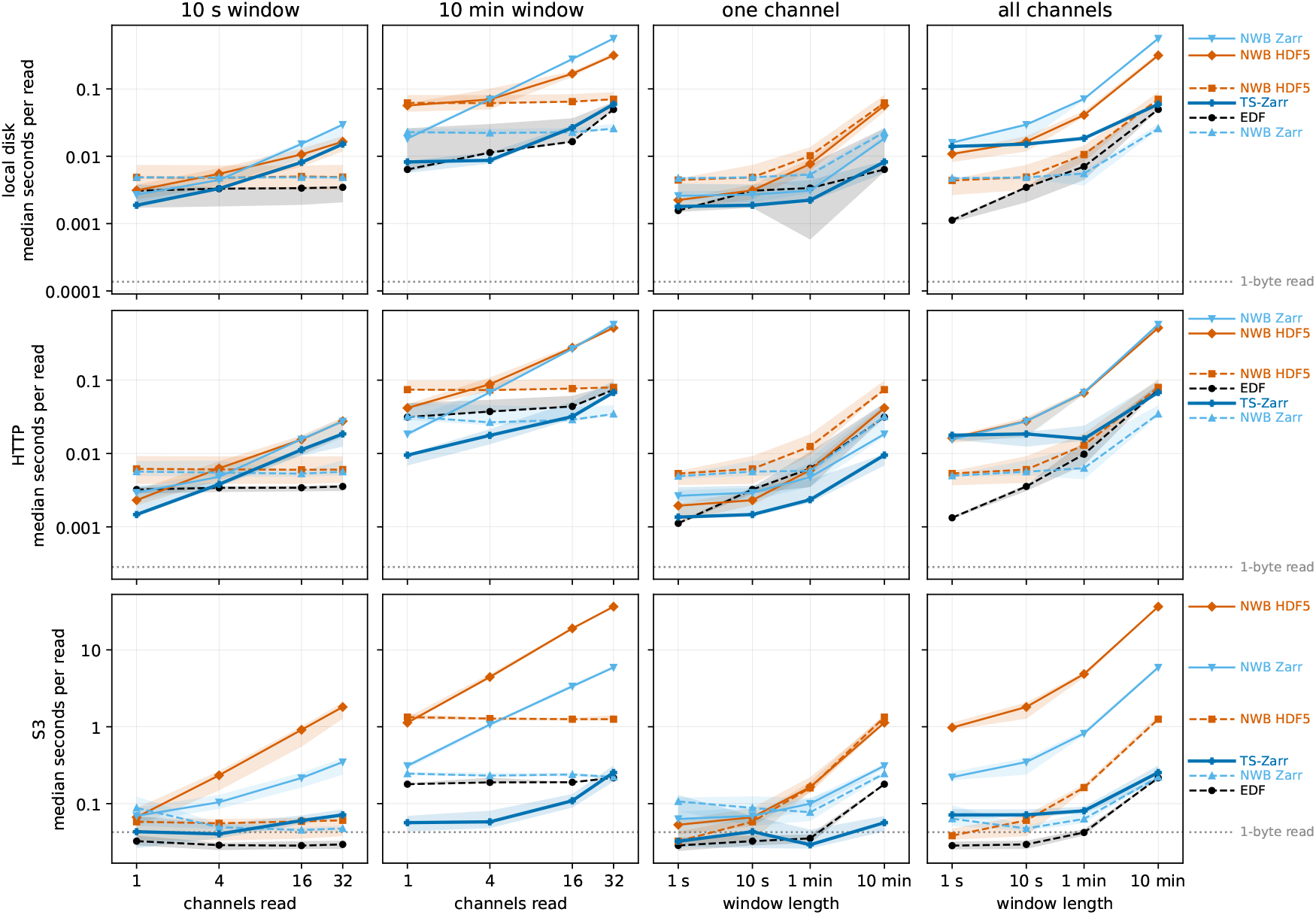
Median seconds per read of the raw samples of the first hour of R2 on local disk, behind HTTP, and on S3, on log axes. The channel panels read windows of 10 s and 10 min from a growing number of channels. The window panels read one channel and all channels at growing window lengths. Solid lines are formats that store each channel separately (TS-Zarr and the NWB files chunked by channel). Dashed lines are formats that store a span of time of every channel together (EDF and the NWB files chunked along time). A TS-Zarr read fetches the raw arrays and no level. Bands span the 25th to 75th percentile of the reads, and the dotted line is the median of a 1-byte read from the same store. The medians cover 5,280 reads on local disk, 5,280 on HTTP, and 4,973 on S3. Each combination of format, read, and window has between one and 100 reads. It stops after the read that brings its total read time to at least 60 s.

**Figure S6:**
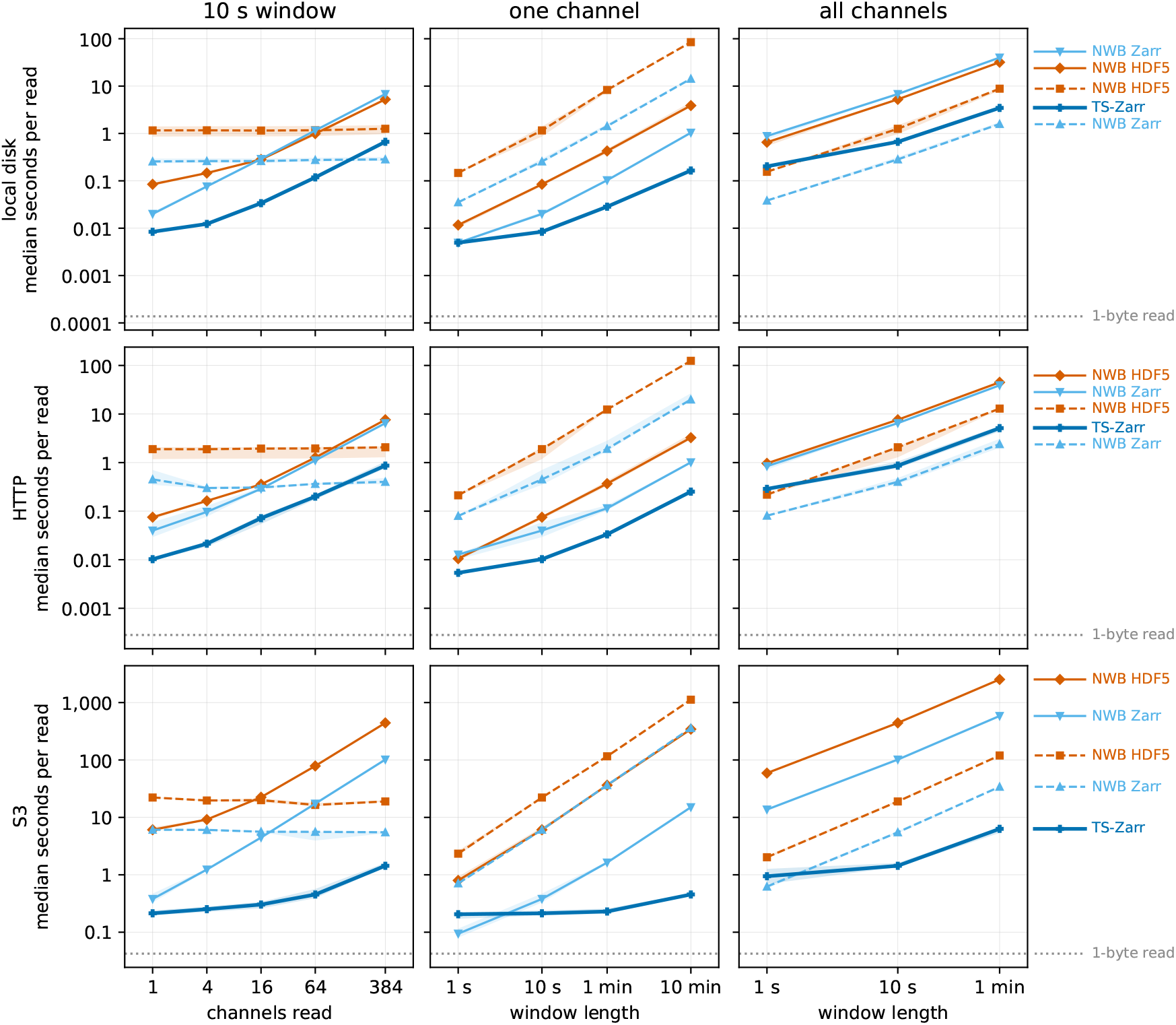
Median seconds per read of the raw samples of R4 on local disk, behind HTTP, and on S3, on log axes. The channel panel reads the 10 s window from a growing number of channels, and R4 has no EDF file. The all-channel read at 10 min is more than 8 GB and is not measured. Solid lines are formats that store each channel separately (TS-Zarr and the NWB files chunked by channel). Dashed lines are formats that store a span of time of every channel together (EDF and the NWB files chunked along time). A TS-Zarr read fetches the raw arrays and no level. Bands span the 25th to 75th percentile of the reads, and the dotted line is the median of a 1-byte read from the same store. The medians cover 3,432 reads on local disk, 3,248 on HTTP, and 1,470 on S3. Each combination of format, read, and window has between one and 100 reads. It stops after the read that brings its total read time to at least 60 s.

**Figure S7:**
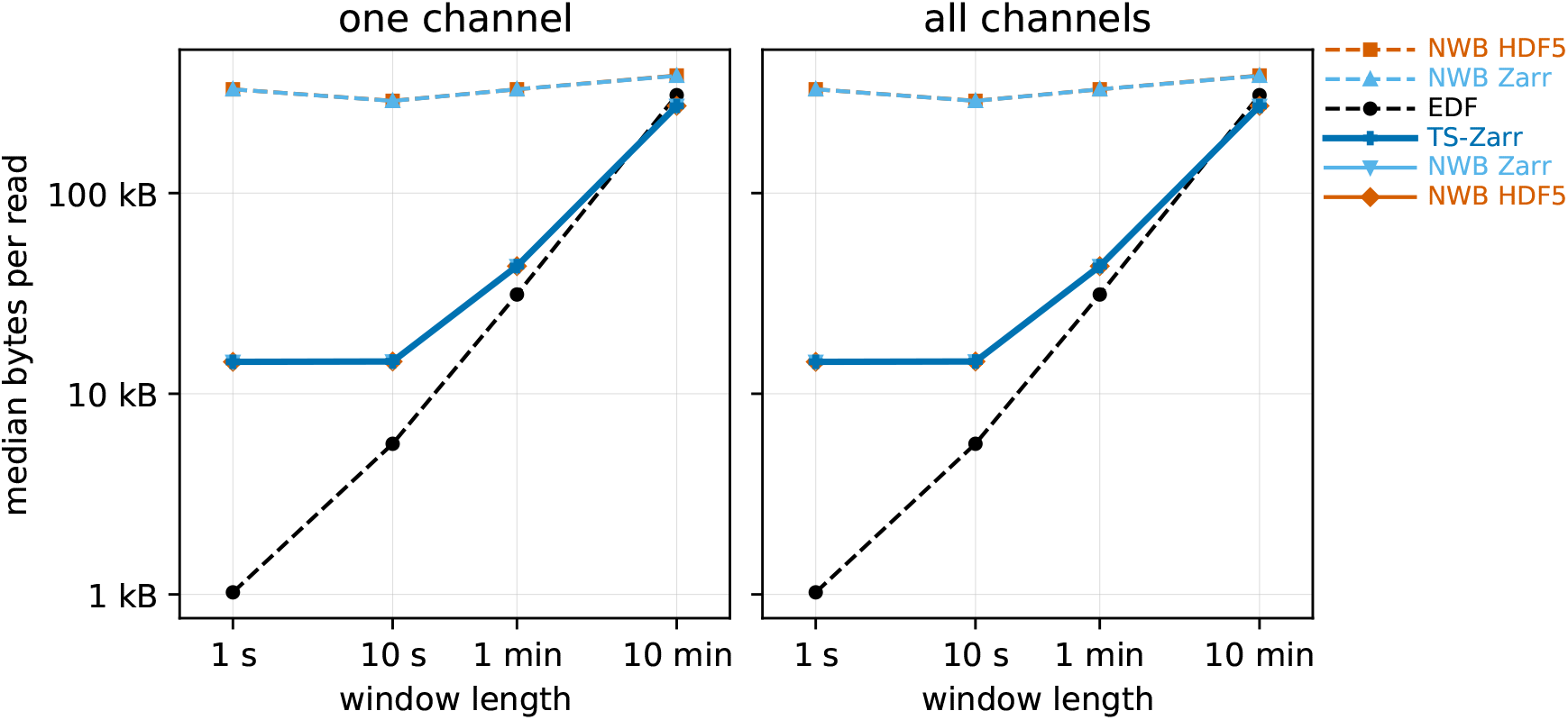
Median bytes that S3 served per read of the raw samples of the first hour of R1, on log axes. The medians cover 3,840 reads. Solid lines are formats that store each channel separately (TS-Zarr and the NWB files chunked by channel). Dashed lines are formats that store a span of time of every channel together (EDF and the NWB files chunked along time). A TS-Zarr read fetches the raw arrays and no level. Each combination of format, read, and window has between one and 100 reads. It stops after the read that brings its total read time to at least 60 s.

**Figure S8:**
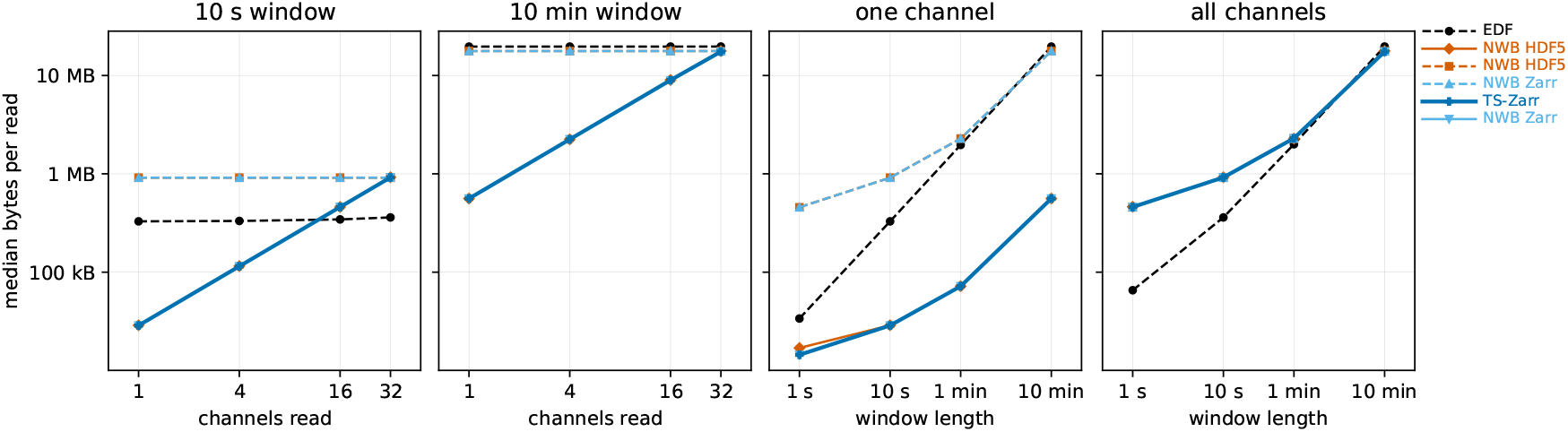
Median bytes that S3 served per read of the raw samples of the first hour of R2, on log axes. The channel panels read windows of 10 s and 10 min from a growing number of channels. The window panels read one channel and all channels at growing window lengths. The medians cover 4,973 reads. Solid lines are formats that store each channel separately (TS-Zarr and the NWB files chunked by channel). Dashed lines are formats that store a span of time of every channel together (EDF and the NWB files chunked along time). A TS-Zarr read fetches the raw arrays and no level. Each combination of format, read, and window has between one and 100 reads. It stops after the read that brings its total read time to at least 60 s.

**Figure S9:**
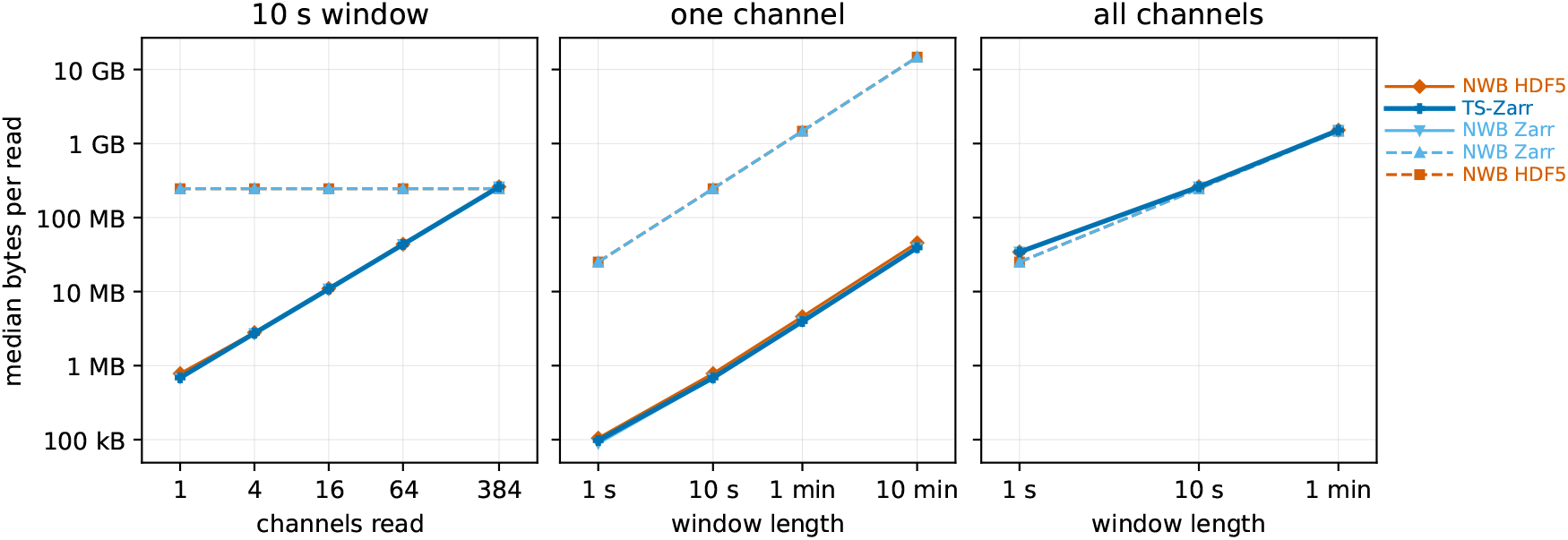
Median bytes that S3 served per read of the raw samples of R4, on log axes. The medians cover 1,470 reads. Solid lines are formats that store each channel separately (TS-Zarr and the NWB files chunked by channel). Dashed lines are formats that store a span of time of every channel together (EDF and the NWB files chunked along time). A TS-Zarr read fetches the raw arrays and no level. Each combination of format, read, and window has between one and 100 reads. It stops after the read that brings its total read time to at least 60 s.

### Resampling benchmark

Each request of the resampling benchmark asked for the `minmax` of every channel over a window at a random start, downsampled to 2000 pixel columns. Windows were 10 s, 1 min, 10 min, 1 h, and 24 h. R4 stopped at its length of 10 min. Both methods downsampled to columns with the `MinMaxDownsampler` of tsdownsample [46]. The resampling server split each channel into groups of whole columns, each spanning about one shard. It read and downsampled the groups on a process pool with one process per vCPU. Each pool process ran up to 32 groups concurrently. For the precomputed read, the load generator picked the coarsest level with at least 2000 bins in the window and fetched its `minmax` alone with 32 threads.

Simulated users ran in closed loop on up to 20 load generators. Each user sent a warm-up request, which was dropped, and 10 measured requests, and stopped 5 min after the start of its run. Requests that took more than 180 s or returned an error were failures and counted as 180 s. Each load generator ran at most 100 users of the server and at most 5 users of the precomputed read. The limit of 5 was the largest number whose CPU use stayed under half of the generator and whose median stayed within 10% of the 1-user median. The precomputed runs at 1000 users exceeded it and were marked client limited. R3 ran 1, 10, 100, and 1000 users at every window, R4 ran 1 and 100, and R1 and R2 ran 1. Once the median request of a run failed, the benchmark skipped the larger user counts of that method and window. It also skipped three server runs of R3. For each window and recording, the benchmark compared the arrays of the two methods once, with NaN equal to NaN, on a window centered on the longest planted gap that fit in it. The arrays were equal in all 18 windows of R1 to R4.

**Table S3:** Median seconds per request of the server divided by that of the precomputed read, on R3. Both methods read from S3. The medians cover 53,060 requests, warm-up requests left out. A cell shows fail where the server run had failures, and not run where a run with fewer users had a median request that failed. A cell ends with (client) where the load generators ran more users than their calibrated limit.

| window | 1 user | 10 users | 100 users | 1000 users |
|---|---|---|---|---|
| 10 s | 0.36 | 0.69 | 5.4 | 17 (client) |
| 1 min | 0.27 | 0.71 | 9.6 | 33 (client) |
| 10 min | 1.2 | 6.3 | 66 | fail (client) |
| 1 h | 5.0 | 37 | fail | not run |
| 24 h | 120 | fail | not run | not run |

**Figure S10:**
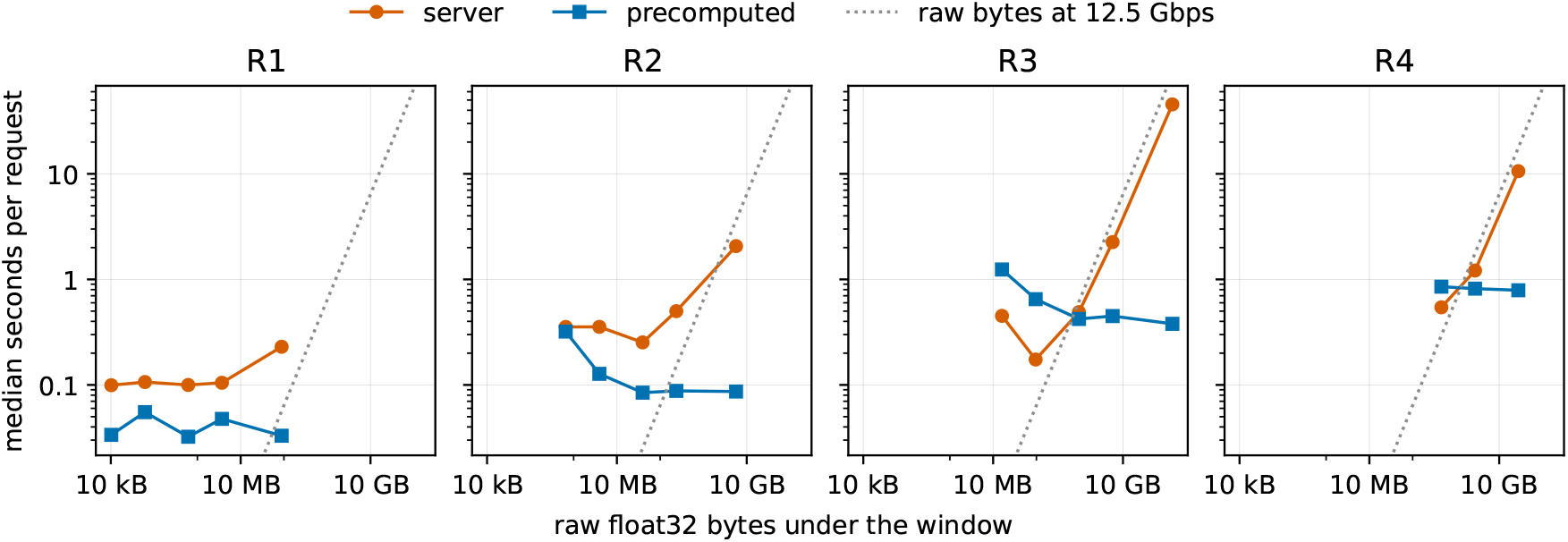
Median seconds per request at one user against the float32 bytes of every channel under the window, for R1 to R4, on log axes. Each panel is one recording, and the panels share both axes. Each line is one method, with one point per window. A request returns the minimum and maximum of every channel over the window in 2000 pixel columns. The dotted line is the time to move the float32 bytes at 12.5 Gbps. Both methods read from S3. The runs cover 356 requests at one user, warm-up requests left out.

**Figure S11:**
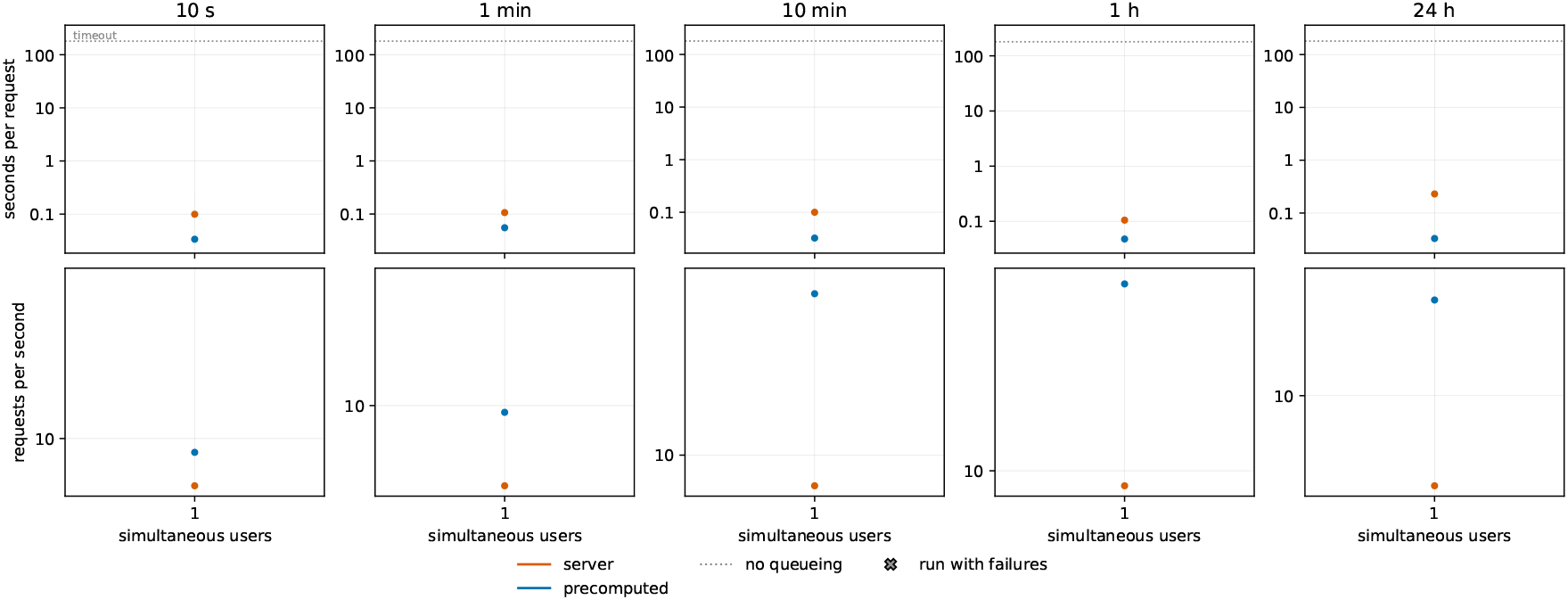
The resampling benchmark on R1 at one user. A request asks for the minimum and maximum of every channel over one window, downsampled to 2000 pixel columns. The upper panels show the seconds per request, as the median and a band from the 5th to the 95th percentile, against the number of users at once. The lower panels show the requests completed per second, with dotted lines for the requests per second if every request took the median of the 1-user run. A cross marks a run with failures, and each failed request counts as 180 s. Both methods read from S3. The runs cover 50 requests of each method, warm-up requests left out.

**Figure S12:**
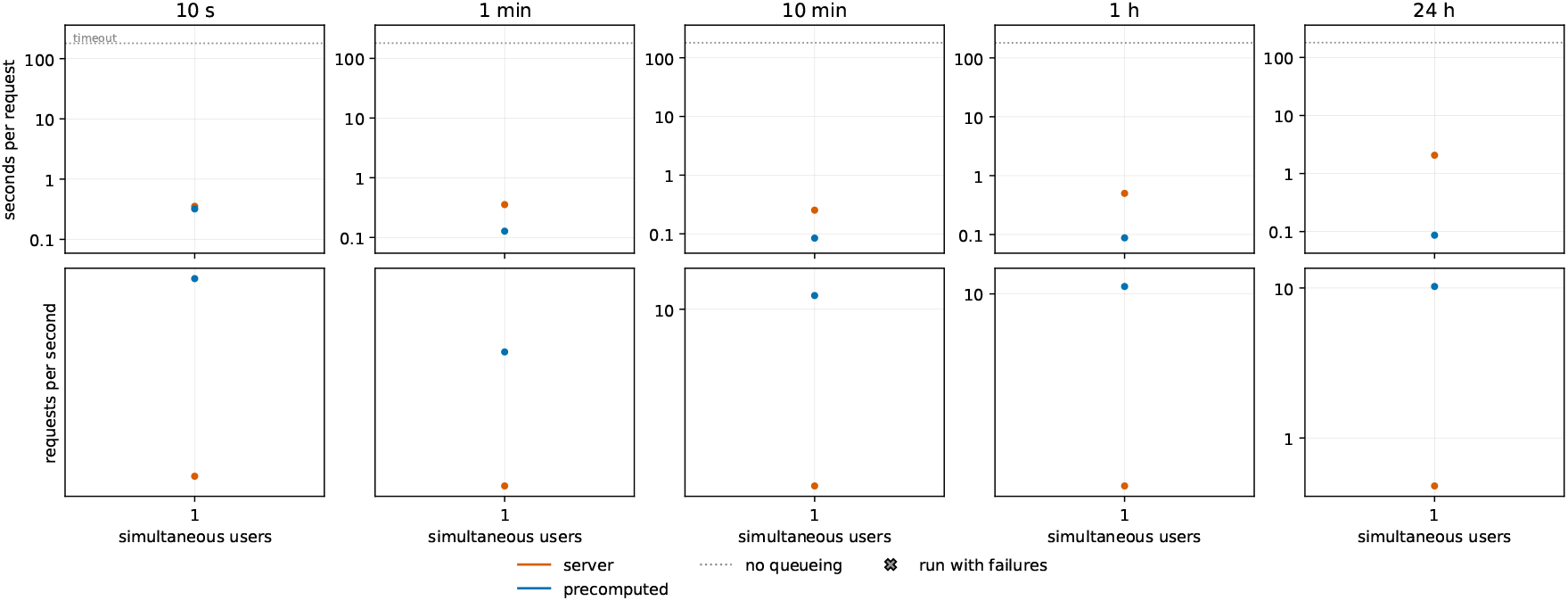
The resampling benchmark on R2 at one user. A request asks for the minimum and maximum of every channel over one window, downsampled to 2000 pixel columns. The upper panels show the seconds per request, as the median and a band from the 5th to the 95th percentile, against the number of users at once. The lower panels show the requests completed per second, with dotted lines for the requests per second if every request took the median of the 1-user run. A cross marks a run with failures, and each failed request counts as 180 s. Both methods read from S3. The runs cover 50 requests of each method, warm-up requests left out.

**Figure S13:**
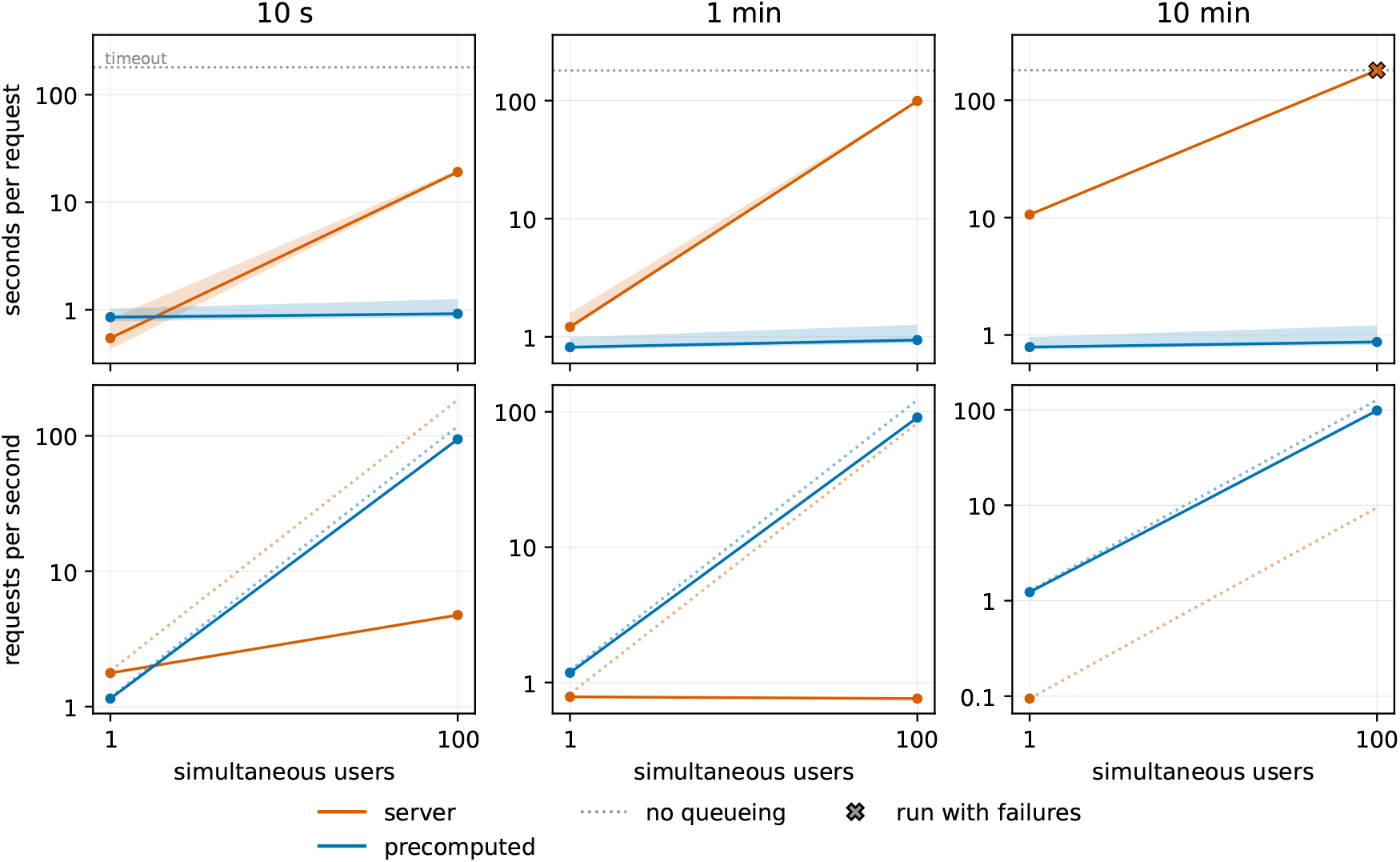
The resampling benchmark on R4 at 1 and 100 users. A request asks for the minimum and maximum of every channel over one window, downsampled to 2000 pixel columns. The upper panels show the seconds per request, as the median and a band from the 5th to the 95th percentile, against the number of users at once. The lower panels show the requests completed per second, with dotted lines for the requests per second if every request took the median of the 1-user run. A cross marks a run with failures, and each failed request counts as 180 s. Both methods read from S3. The runs cover 1,438 server requests and 3,030 precomputed requests, warm-up requests left out.

